# Tissue topography steers migrating Drosophila border cells

**DOI:** 10.1101/2020.09.27.316117

**Authors:** Wei Dai, Xiaoran Guo, Yuansheng Cao, James A. Mondo, Joseph P. Campanale, Brandon J. Montell, Haley Burrous, Sebastian Streichan, Nir Gov, Wouter Jan Rappel, Denise J. Montell

## Abstract

Moving cells can sense and respond to physical features of the microenvironment, however *in vivo* the significance of tissue topography is mostly unknown. Here we use the Drosophila border cells, an established model for *in vivo* cell migration, to study how chemical and physical information influence migration path selection. Live imaging, genetics, modeling, and simulations show that, although chemical cues were thought to be sufficient, microtopography is also important. Chemoattractants promote predominantly posterior movement, whereas tissue architecture presents orthogonal information, a path of least resistance concentrated near the center of the egg chamber. E-cadherin supplies a permissive haptotactic cue. Our results provide insight into how cells integrate and prioritize topographical, adhesive, and chemoattractant cues to choose one path amongst many.

## Introduction

Cell migrations are essential in development, homeostasis, and disease. While chemoattractants and repellents have been extensively studied (1–3), physical features of the microenvironment may be equally important. Here we use Drosophila border cells as a model and uncover a role for tissue topography in directional cell migration *in vivo*. Border cells are 6-10 follicle cells that delaminate and migrate collectively ~150 μm over 3-6 hours within ovarian egg chambers, which are composed of 15 nurse cells and one oocyte, encased within ~850 epithelial follicle cells (4–6) (Fig.1A, movie 1).

**Fig. 1.**
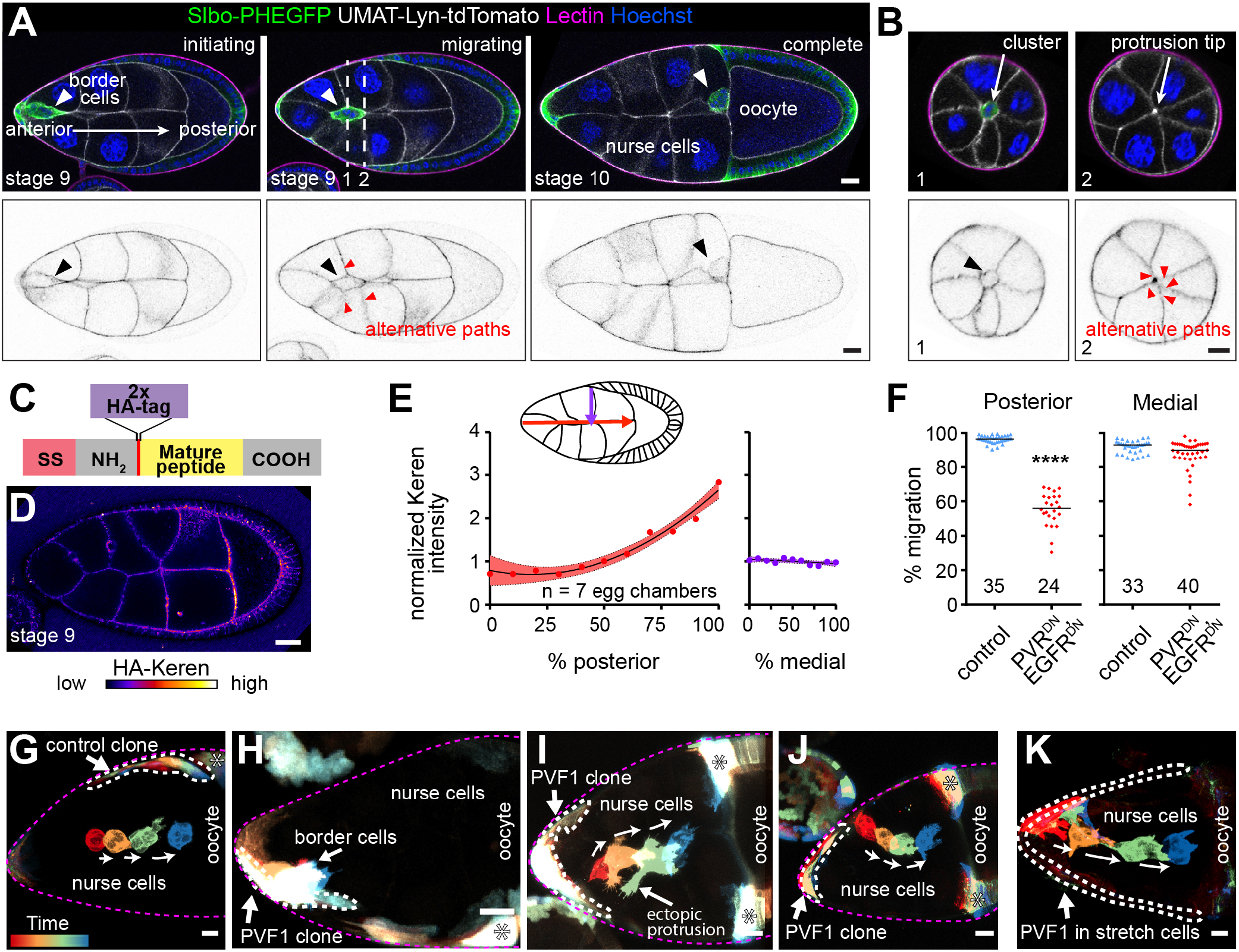
Medial migration is not primarily controlled by chemoattractant guidance cues. (A) Lateral views of egg chambers showing border cell migration between nurse cells to the oocyte. Dashed lines represent cross sections shown in (B). (C) Schematic of endogenously CRISPR-tagged HAKeren. (D) A living, unpermeabilized egg chamber stained for anti-HA-Keren (see also fig. S2). (E) Quantification of HA-Keren along anterio/posterior (left) and mediolateral (right) axes. Each dot represents a location along the path. (F) Quantification of border cell position. Each dot, one cluster. ****, P < 0.0001 (Mann-Whitney test). (G to K) Rainbow views of border cell migration in control or with ectopic UAS-PVF1 expression in clones (H-J) or in all anterior, “stretch” follicle cells (K). Scale bars, 20 μm.

The oocyte secretes chemoattractants that activate two receptor tyrosine kinases (RTKs) (7–9). The platelet derived growth factor and vascular endothelial derived growth factor like protein (PVF1) activates its receptor PVR (9). The ligands Spitz (Spi), Keren (Krn), and Gurken (Grk) activate the Drosophila epidermal growth factor receptor (EGFR) (10). Border cells lacking expression or activity of both RTKs, fail to reach the oocyte (8, 9), and ectopic PVF1, Spi or Krn is sufficient to reroute them (10). Similarly, lymphocyte homing, axon pathfinding, and migration of the zebrafish lateral line (11), neural crest (12), and primordial germ cells (13) have been attributed primarily to chemoattraction/repulsion. While the effects of substrate stiffness on migrating cells have been studied in vitro and *in vivo* (14–16), other physical features, like tissue topography, remain relatively unexplored.

By reconstructing egg chambers in three dimensions (3D), we noticed two orthogonal components to border cell pathway selection. Border cells migrate from anterior to posterior, the obvious path in a typical lateral view (Fig. 1A, fig. S1A). In addition, they follow a central path (Fig. 1B, fig. S1B and C, movie 1), despite encountering ~40 lateral alternatives (Fig. 1B and movie 2).

## Results

To address whether the extracellular RTK ligands are present in gradients that might explain both posterior and medial guidance, we used CRISPR to epitope-tag endogenous PVF1, Spi, and Krn (see methods). [Grk directs dorsal movement only as the cells near the oocyte (4)]. Extracellular HA-tagged Krn (Fig. 1C) accumulated in an anterior (low) to posterior (high) gradient; however its concentration was not higher medially than laterally (Fig. 1D and E, fig. S2A and B). Intracellular, but not extracellular, HA-tagged PVF1 was detectable (fig. S2C and D). Tagged Spi was undetectable.

Since we could not detect all of the ligands, we addressed their contributions by expressing dominant-negative receptors (PVRDN and EGFRDN), which impedes posterior migration (8) (Fig. 1F, fig. S3A and B). Importantly, mediolateral defects were rare, occurring in <10% of egg chambers (Fig. 1F). RNAi caused similar effects (fig. S3C). Therefore, some other factor(s) must guide the cells medially.

Live imaging of egg chambers with ectopic PVF1 provided further evidence for independent attraction to the egg-chamber center (Fig. 1G to K). When anterior follicle cells ectopically expressed PVF1, border cells sensed the ligand because they were frequently detained at the anterior (Fig. 1H) compared to controls (Fig. 1G). Border cells also frequently protruded toward the ligand-expressing cells but remained on the central path (Fig. 1I). In other cases (Fig. 1J), border cells migrated along a patch of PVF1-expressing follicle cells, lingered, then nevertheless left the clone and returned to the egg-chamber center, ignoring more direct routes to the oocyte. PVF1 expression in all anterior follicle cells produced similar results (Fig. 1K). Thus even in the presence of ectopic chemoattractant, border cells preferred the egg-chamber center, again suggesting that another signal steers them medially.

Of all the migration-defective mutants analyzed, only nurse-cell knockdown of E-cadherin exhibits dramatic mediolateral defects (17) (Fig. 2A and B), causing border cells to move between E-cadherin-positive follicle cells and nurse cells (fig. S4, movie 3).

**Fig. 2.**
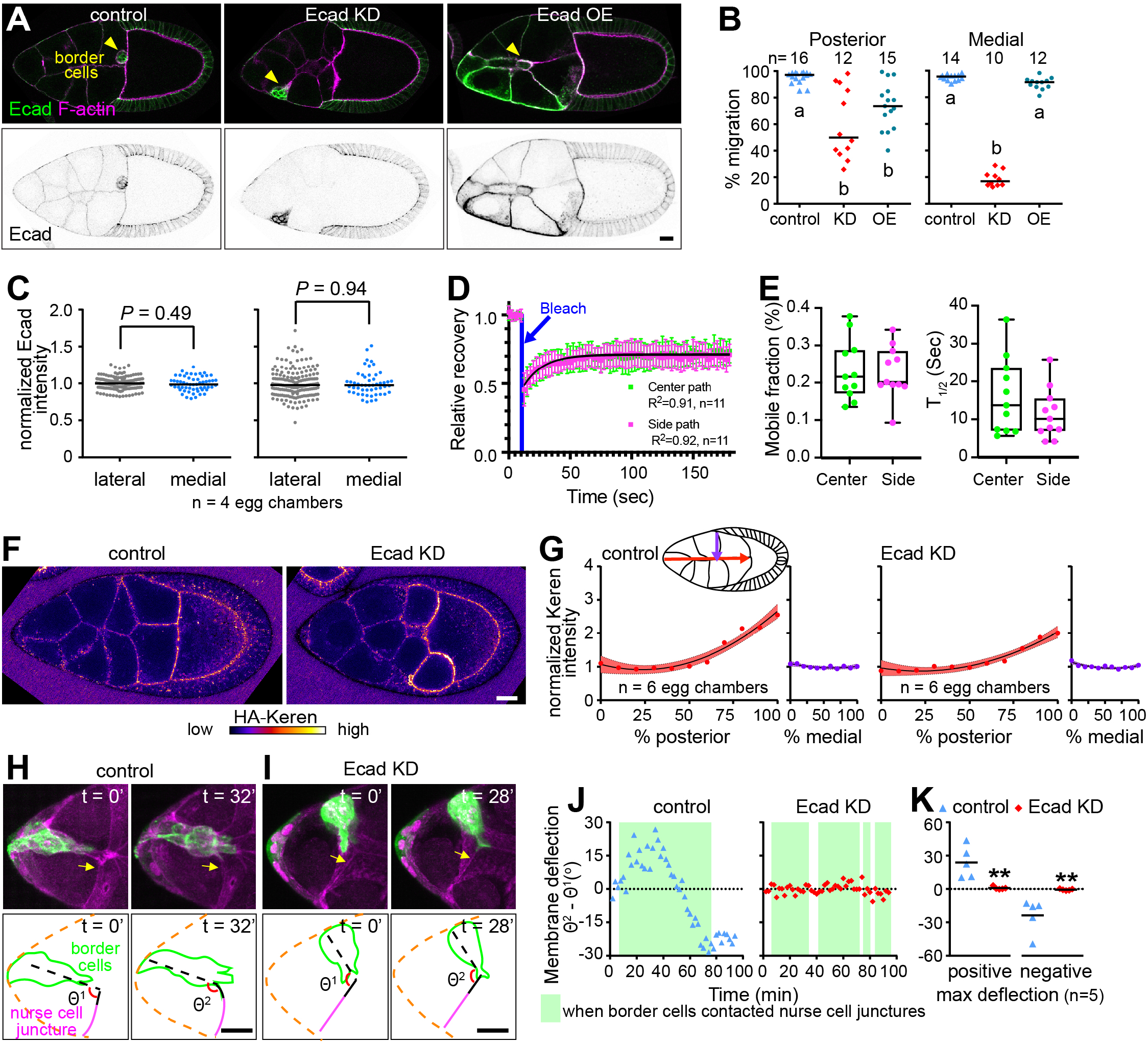
E-cadherin, a permissive medial traction cue. (A) Images of egg chambers from control, nurse-cell E-cadherin (Ecad) knockdown (KD), or mosaic nurse-cell overexpression (OE). Top panels show anti-E-cadherin (green) and phalloidin staining of F-actin (magenta). Bottom panels show anti-E-cadherin alone. (B) Quantification of migration. Letters a and b designate significantly different groups (*P* < 0.01, Kruskal-Wallist test). (C) Quantification of E-cadherin and F-actin intensities on medial and lateral surfaces that lack ring canals. Anova with post-hoc Tukey test and blocking by egg chamber. (D) Quantification of the FRAP experiment shown in (fig. S6). (E) Quantification of the mobile fraction and T1/2 of E-cadherin on center and side membranes. (F) Anti-HA-stained, living egg chambers from control and E-cadherin knockdown (KD). (G) Quantification of Keren intensity along the anterior-posterior (coral) and lateral-medial (purple) axes. The line represents the best fit trendline of normalized Keren distribution along the A/P or L/M axis. Shading represents the standard deviation of the best fit trendline. There is no statistically significant difference between control and nurse-cell Ecad RNAi along the mediolateral axis (*P* = 0.6) or the relevant portion of the A/P axis (*P* = 0.6) that border cells encounter in the nurse-cell Ecad RNAi condition (see Fig. 2B). (H and I) Still images from movies showing (H) border cells pull on and deflect nurse-cell junctures in the control but not in the E-cadherin knockdown (I). (J) Traces of nurse-cell membrane deflections. (K) Quantification of maximum deflections. (**, *P* < 0.01 Mann-Whitney test). Scale bars, 20 μm.

How does nurse-cell E-cadherin contribute to central path selection? Neither uniform (17), nor asymmetric E-cadherin overexpression on nurse cells caused any medial guidance defect (Fig. 2A and B). Using near isotropic light sheet imaging (fig. S5), we detected no significant difference in E-cadherin concentration (Fig. 2C) on central versus side paths. Fluorescence recovery after photobleaching (fig. S6) also revealed no difference in E-cadherin dynamics between central and side paths (Fig. 2D and E). E-cadherin knockdown did not alter the HA-Krn distribution in a way that could account for lateral path selection (fig. S7): a mediolateral gradient was absent, and an anterior/posterior gradient was still present (Fig. 2F and G).

Follicle cells normally express more E-cadherin than nurse cells (fig. S8A), so if differences in E-cadherin concentration steered border cells, we would expect that reduction of follicle cell E-cadherin might cause mediolateral guidance defects; yet follicle cell RNAi caused no defect (fig. S8B). Nor did E-cadherin over-expression in follicle cells impact pathfinding (fig. S8C to E). Therefore, though the presence of a low level of E-cadherin normally found on nurse cells is essential for border cells to migrate between them, we found no evidence that E-cadherin concentration differences were sufficient to steer border cells.

We noticed that border cells pulled on wild-type nurse-cell membranes as they migrated, creating a measurable deflection of the nurse cells (Fig. 2H, movie 4, fig. S9). In contrast, border cells protruding between E-cadherin-negative nurse cells did not deflect their membranes (Fig. 2I to K, movie 4), suggesting border cells could not get traction. The lack of traction could in principle fully account for their inability to take the central path between nurse cells. Interestingly, border cells still protruded extensively between nurse cells; however the protrusions appeared as broad flat lamellipodia (movie 3) rather than the normal spear-shaped protrusions (movie 1). We conclude that E-cadherin supplies a permissive traction cue. As previously described (17), this mechanical function amplifies RTK signaling and shapes forward protrusions (17 and this study); however something other than differential adhesion must normally steer border cells to the central path.

Since neither chemoattractant nor adhesive cues fully accounted for medial pathfinding, we reconstructed egg chambers in 3D and characterized central versus side migration paths. The nurse-cell-oocyte complex is a syncytium packed within the follicular epithelium (fig. S10, movie 8) (18). A striking feature of the central path is that it is where ≥3 nurse cells come together (lines and magenta dots in Fig. 3A, fig. S11). Side paths are largely composed of 2-nurse-cell interfaces (lines in Fig. 3B; planes in Fig. 3C, movie 5). Strikingly, ≥3-nurse-cell junctures are enriched near the center (Fig. 3D). We use the word juncture, rather than junction to reflect the fact that they are places where cells come together but are not adherens, tight, or gap junctions.

**Fig. 3.**
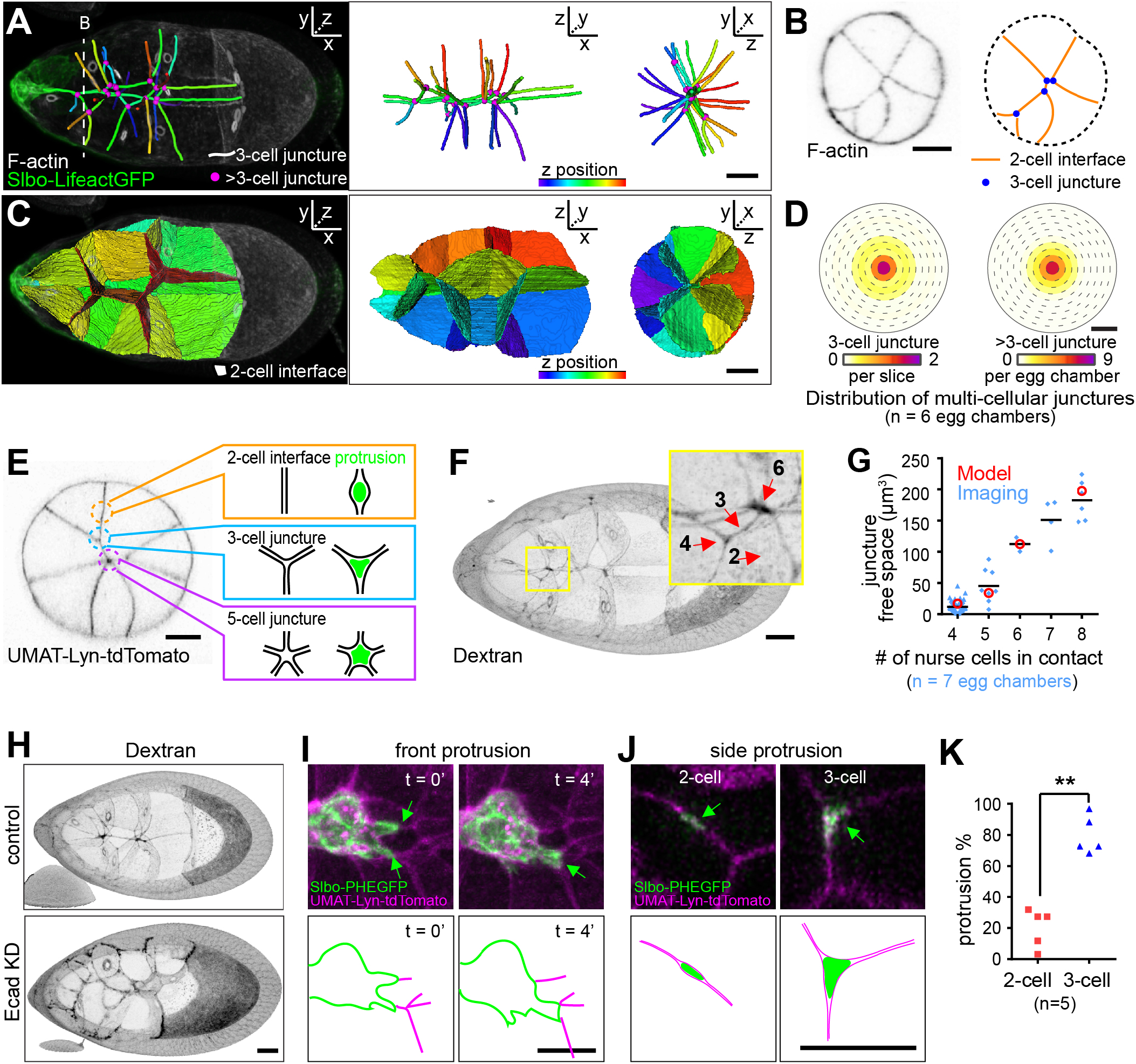
The central path is enriched in junctures where multiple nurse cells meet. (A to C) 3D reconstructions of nurse-cell contacts. Dashed line in (A) indicates cross section in (B). (D) Heat map showing distributions of 3 (left) or >3 (right) cell junctures as a function of mediolateral position. (E) Schematic representation of border cell protrusion into nurse-cell junctures illustrating the concept that more space is expected in regions where more nurse cells meet. (F) Extracellular spaces filled with fluorescent dextran in a wild type egg chamber. (G) Quantification of extracellular volume as a function of the number of cells in a juncture. Values from the 3D model (red) (supplementary text ST1) and the experimental data (blue). (H) Dextran filled spaces (black) in a control stage 9 egg chamber compared to germline E-cadherin knockdown (KD). (I) Still images from two time points from a 3D movie. Two forward-directed protrusions (green arrows in I) encounter different multiple-nurse-cell junctures (magenta). The protrusion that reaches a >3 cell juncture wins (green arrow in B, n = 11 from 7 movies). (J) Cross-sections showing side protrusions (green) at 2-cell interface or 3-cell junctures (magenta). (K) Percentage of total side protrusions extending into 2-cell or 3-cell junctures. **, *P* < 0.01 (paired t test). Scale bars, 20 μm.

We considered the influences that this geometry would likely have on border cells squeezing between nurse cells (supplementary texts ST1 and ST2). Due to the energy cost of unzipping nurse-cell-nurse-cell adhesions, protrusion into regions where multiple nurse cells meet should in principle be more favorable (Fig. 3E). This geometry argument predicts that there should be larger spaces where more nurse cells come together (fig. S12A to D), which we confirmed by measuring extracellular spaces using fluorescent dextran (Fig. 3F and G). As predicted, germline E-cadherin knockdown opened larger spaces (Fig. 3H, fig. S12E), confirming that E-cadherin normally seals nurse cells together. The free space should be most relevant at the scale of protrusions similar in size to the open space; the protrusions then steer the cluster. In vitro, migrating cells have been shown to choose channels that accommodate the size of the nucleus (19, 20); here we show that *in vivo,* even smaller spaces can guide cells.

To test the prediction that crevices where more cells come together present a lower energy barrier for protrusion, we examined 3D movies. Junctures of ≥3 nurse cells line the center path, and forward-directed protrusions always extended between multiple nurse cells. Moreover, when cells encountered two ≥3-nurse-cell paths, the cluster extended two protrusions (Fig. 3I). Eventually, the protrusion between the greater number of nurse cells always won.

When cells probed side paths, they extended into both 2-cell and 3cell junctures (Fig. 3J). However, protrusion into three-nurse-cell junctures were more frequent (Fig. 3K), even though 2-nurse-cell interfaces offer vastly greater surface area (Fig. 3A and C). We conclude that crevices where multiple nurse cells come together create an energetically favorable path, and tissue topography, specifically ≥3-cell junctures, normally promotes central pathway selection.

To test whether the combination of an anteroposterior chemoattractant gradient and a bias toward multiple cell junctures is in principle sufficient to explain border cell behavior, we developed a dynamic model that describes the trajectory of border cells moving within a realistic egg chamber geometry (Fig. 4A). We modeled the border cell cluster as a particle that moves stochastically in an effective potential 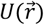 (ST3) that incorporates two independent guidance terms: 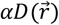, the energy cost for the cluster to move between N nurse cells, and 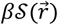, the anteroposterior chemoattractant gradient. Simulating normal border cell migration conditions replicated normal trajectories (Fig. 4A to C, and movie 6). Eliminating the chemoattractant caused significant posterior migration defects but little deviation from the central path (Fig. 4A to D), consistent with experiments (Fig. 1F). In contrast, eliminating the preference for ≥ 3-nurse-cell junctures randomized mediolateral path selection without posterior migration defects (Fig. 4A, B, and E). Eliminating both terms produced dramatic mediolateral and anterior-posterior defects (Fig. 4A, B, and F).

**Fig. 4.**
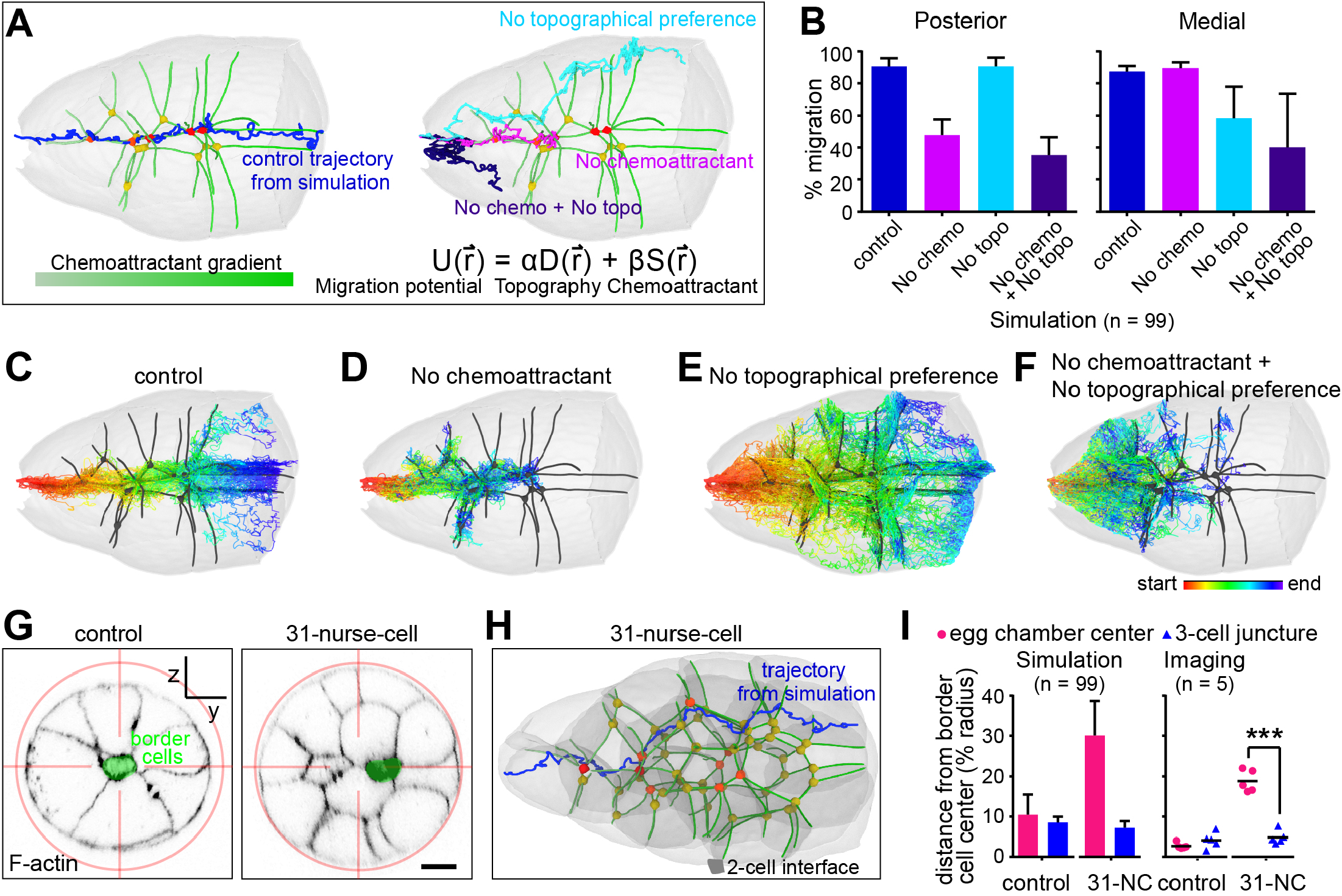
Simulations and experiments showing preference for multiple cell junctures over the geometric center. (A) Representative simulated trajectories through the wild-type geometry shown in Fig. 3A. (B) Quantification based on 99 simulations. (C to F) Rainbow display of 99 simulated migration paths (red=start and blue=end) for each of the indicated conditions. (C) Normal condition in which path selection is a function of both a posterior chemoattractant gradient and increasing preference for junctures with increasing numbers of nurse cells. At the end of migration the influence of geometry weakens, correlating with loss of >3 nurse cell junctures. (D) Removal of the posterior chemoattractant gradient causes 100% posterior migration defect and 10% medial migration defect. Those clusters that deviate from the central path migrate into side paths with >3-cell junctures. (E) In the presence of chemoattractant but absence of preference for multiple nurse cell junctures, clusters migrate posteriorly but are distributed essentially randomly with respect to the mediolateral axis. (F) In the absence of both chemoattractant and junctional preference, clusters exhibit both posterior and medial migration defects. (G) Cross-sections showing border cell and nurse-cell positions relative to the egg-chamber center in a control compared to a 31-nurse-cell egg chamber. (H) Representative simulated trajectory. (I) Comparison of the distance from the border cell centroid to the egg-chamber center vs the nearest 3-cell juncture. In both simulations and experiments, border cell position correlates more strongly with 3-cell junctures than the geometric center. ***, *P* < 0.001 (Paired t-test). Scale bars, 20 μm.

Multiple cell junctures are concentrated near the egg chamber center, so it could be that border cells are attracted to multiple cell junctures or to some other property of the geometric center of the egg chamber. For example, it could be that border cells are attracted to the center due to an unknown, centrally-concentrated chemoattractant or possibly due to an unknown laterally-concentrated chemorepellent, or some other unknown factor. To distinguish whether border cells prefer multiple cell junctures or the geometric center, we analyzed egg chambers with atypical geometries. In mutants that alter early germ cell divisions (21), we found some 31-nurse-cell egg chambers (fig. S13) with a central 2-nurse-cell interface (Fig. 4G). In each instance, the border cells selected the ≥3 nurse-cell-junctures even when off-center (Fig. 4G). Simulating migration using the 31-nurse-cell geometry and the same parameters as for wild-type reproduced the result (Fig. 4H and I). These results support the interpretation that border cells are attracted to multiple-cell junctures over any other property of the geometric center.

We also simulated migration in egg chambers lacking nurse-cell E-cadherin, in which there is more free space where two nurse cells meet follicle cells than between one nurse cell and follicle cells (Fig. 5A). The model predicted and experiments confirmed that the border cells zig zag along grooves at nurse cell-nurse cell-follicle cell junctures (Fig. 5B, fig. S14, and movie 7).

**Fig. 5.**
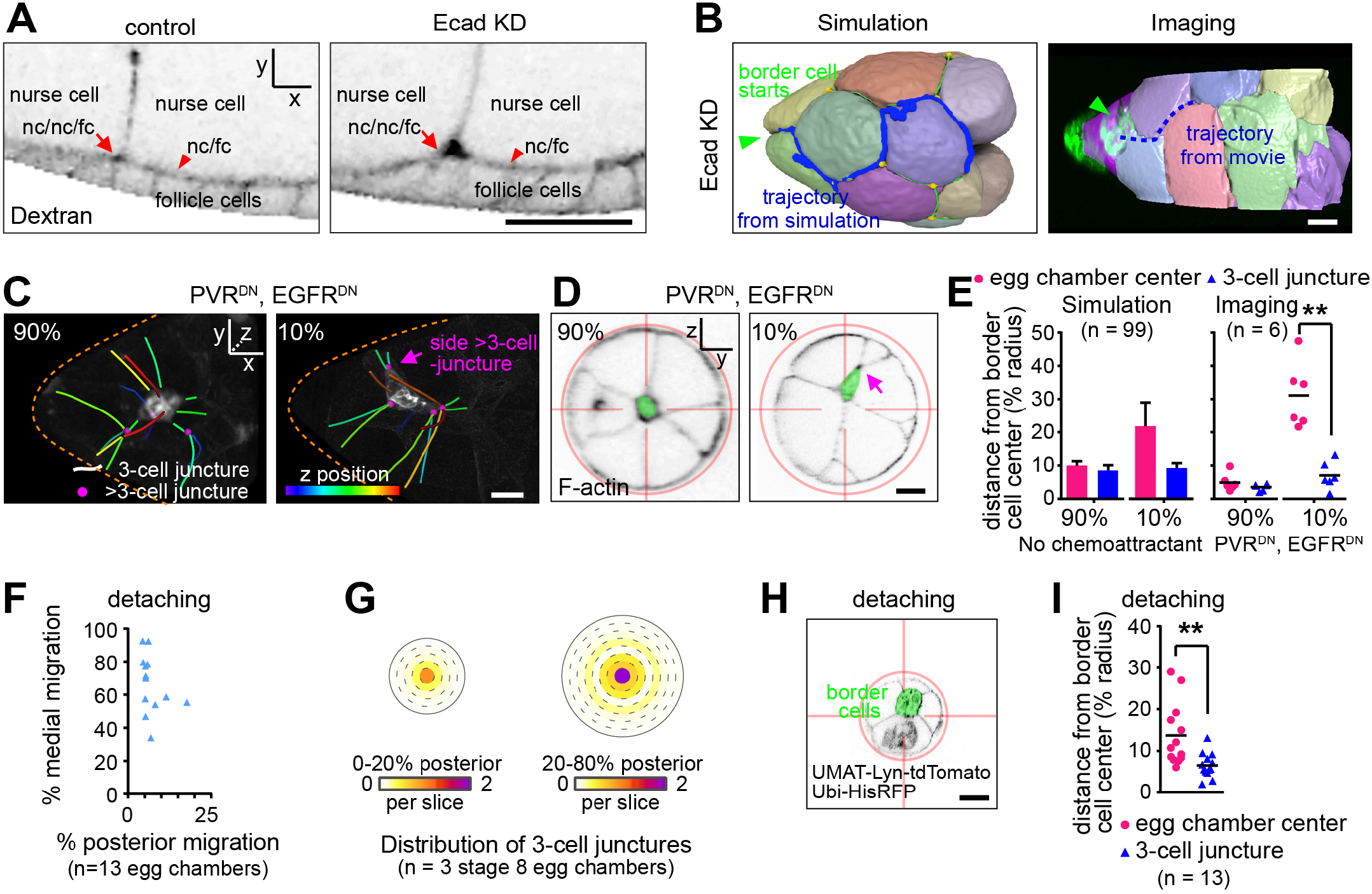
Border cells follow off-center multiple-cell junctures. (A) Lateral view of dextran showing E-cadherin knockdown enlarged space. (B) 3D reconstructions of nurse cells with E-cadherin knockdown showing border cells in nurse cell-nurse cell-follicle cell grooves. (C and D) Border cell cluster location relative to nearby ≥3-cell junctures in representative egg chambers in which border cells express PVRDN and EGFRDN. In 90% of egg-chambers, border cell clusters remain in the center, while 10% migrate off-center (Fig. 1F, right panel). (E) Quantification of distance from border cell centroid to the egg chamber center and the nearest 3-cell juncture. Simulation of no chemoattractant is done by removing the chemoattractant term. **, *P* < 0.01 (Paired t-test). (F) Migration in control stage 9 egg chambers when border cells are detaching from anterior follicle cells. (G) Heat map showing distributions of 3-cell junctures as a function of mediolateral position in stage 8 egg chambers at 0-20% posterior location or 20-80% posterior location. (H) Cross-sections showing border cells (green pseudocolor) and nurse cell junctions (black lines) relative to the egg chamber center. (I) Comparison of the distance from the border cell centroid to the egg chamber center (pink dots) vs the nearest 3-cell junction (blue triangles) in early stage 9 egg chambers when border cells are detaching from anterior follicle cells. **, *P* < 0.01 (Paired t-test). Scale bar, 20 μm.

We then re-examined the 10% of PVRDN, EGFRDN egg chambers in which border cells are found off-center (Fig. 1F). Remarkably, border cells again moved to sites where multiple nurse cells meet (Fig. 5C and D), supporting the idea that multiple-cell junctures are energetically favorable even when not at the geometric center. Simulations also recapitulated this result (Fig. 5E).

At the initiation of migration when border cells first detach from the anterior follicle cells, the border cells are not always located at the geometric center (Fig. 5F to I), and 3-nurse-cell junctures are not as concentrated at the center at the anterior, compared to the rest of the migration path (Fig. 5G). Again, border cells preferred multiple-nurse-cell-junctures over the geometric center (Fig. 5H and I).

Many other features of the central path proved inconsequential (Fig. 6). For example, residual, stabilized cleavage furrows called ring canals connect germline cells to one another in a regular pattern (fig. S10). While the central path typically lacks ring canals (Fig. 6A, top), 63% lateral paths also lack ring canals (Fig. 6A, bottom), so absence of ring canals is insufficient to direct border cells centrally. Occasionally, we observed a ring canal along the migration path (Fig. 6B, fig. S15), and the border cells migrated around it (Fig. 6B). So absence of ring canals is neither necessary nor sufficient to provide medial guidance.

**Fig. 6.**
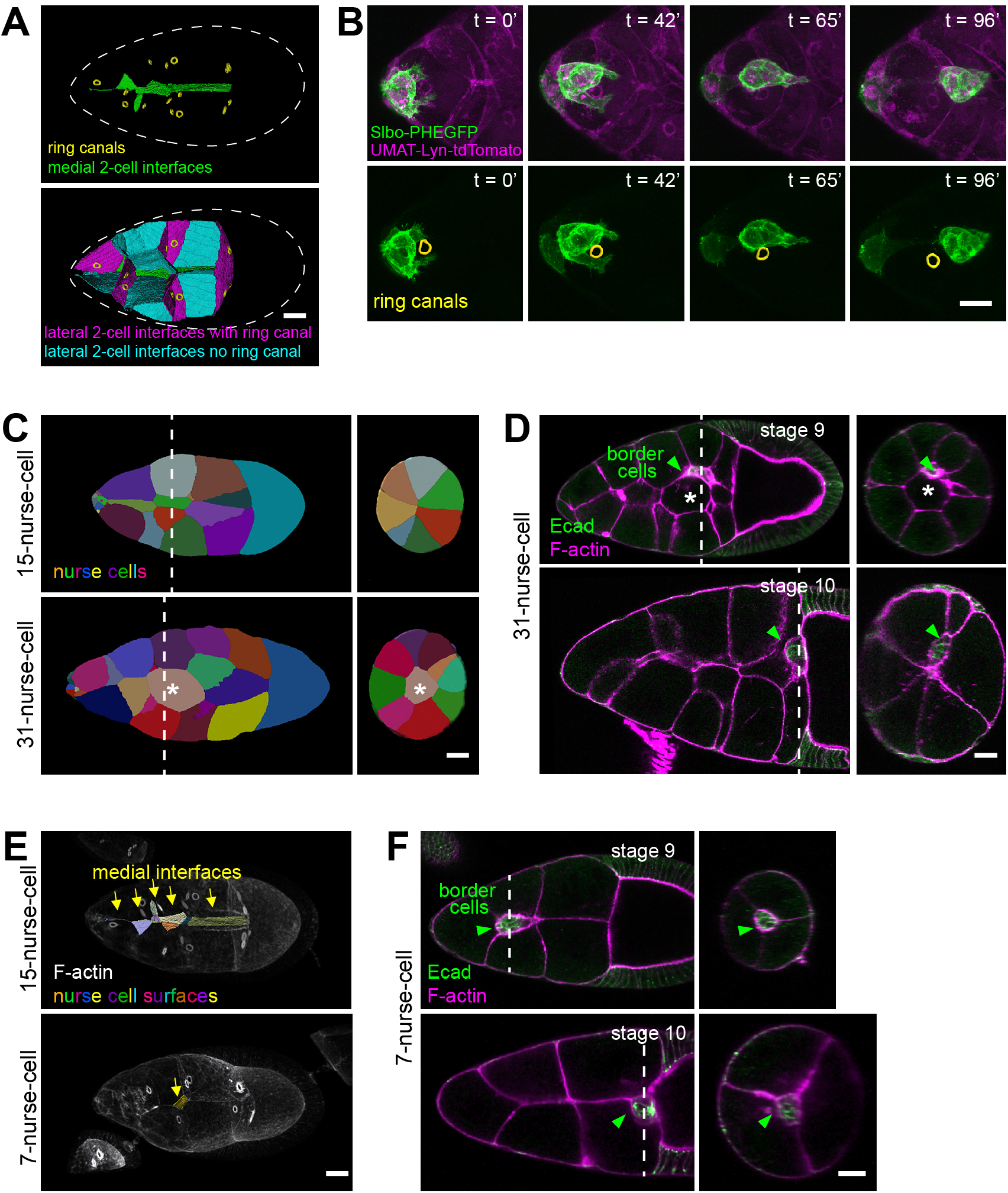
Some physical features are not critical. (A) Ring canal locations on 2-germ cell interfaces. Medial interfaces (green) are free of ring canals (left panel). 15 lateral cell interfaces contain ring canals (magenta, right panel) and 26 do not (cyan). (B) Still images from a time lapse movie showing border cells migrating past a ring canal. Dashed line outlines the egg chamber. (C) Reconstructed nurse cell arrangements showing all nurse cells contact follicle cells in control, but there are 1-2 nurse cells in the center in 31-nurse-cell egg chambers. (D) Border cells migrate around a central nurse cell in stage 9 (n=9) and reach the oocyte border by stage 10 in an egg chamber with 31 nurse cells (n=19). White lines indicate cross-sectional planes shown in the right panels. White asterisks indicate the central nurse cell. (E) Lateral view of segmented 3D nurse cell medial surface-surface contacts in early stage 9 egg chambers. (F) Border cells migrated along the multiple cell juncture and reached the oocyte border in stage 10 egg chambers with 7 nurse cells (n=41). White dashed lines indicate cross sectional plane shown in the right panel. Scale bars, 20 μm.

The central path is also normally devoid of cells. However, in 31 nurse cell egg chambers, border cells successfully navigate around centrally located cells (Fig. 6C and D). The central path is also normally composed of much smaller surface areas compared to side paths (Fig. 6A, fig. S16A and B). However, in egg chambers with 31 nurse cells (fig. S16A and B), the differences between medial and lateral surface areas are much reduced (fig. S16B), and in egg chambers with 7 nurse cells (fig. S16C and D) (22), the medial surfaces are nearly absent (Fig. 6E). Nevertheless, border cells still migrate in multicellular junctures and reach the oocyte border (Fig. 6F).

We gained further insight into how the cells integrate and prioritize the chemical and geometric cues. Results reported here show that, for most of their trajectory, the chemoattractants primarily guide the border cells posteriorly and multicellular junctures steer them centrally. These findings are consistent with an earlier study that used photo-activatable Rac (PA-Rac) to steer border cells (23). PA-Rac could steer cells forward or backward down the center path throughout migration but was only able to steer them off center near the beginning or near the end (23). Since Rac functions downstream of the RTKs, this already suggested that some other cue must steer the cells centrally and that this cue should be strongest between 25% and 75% of the distance to the oocyte.

Interestingly, ≥3 cell junctures near the end of migration, as border cells approach the oocyte, we found that >3-cell junctures are absent (Fig. 3A), which the model predicts would weaken the central bias of topographical information. Chemoattractant levels are highest near the oocyte, and the EGFR ligand Gurken is enriched dorsally (10). By aligning egg chambers according to the position of the oocyte nucleus, we noticed that the border cells typically squeeze between two nurse cells to move dorsally (Fig. 7A and B, movie 9). Adding Grk into the model and simulation accurately predicts this dorsal turn (Fig. 7C and D, movie 10).

**Fig. 7.**
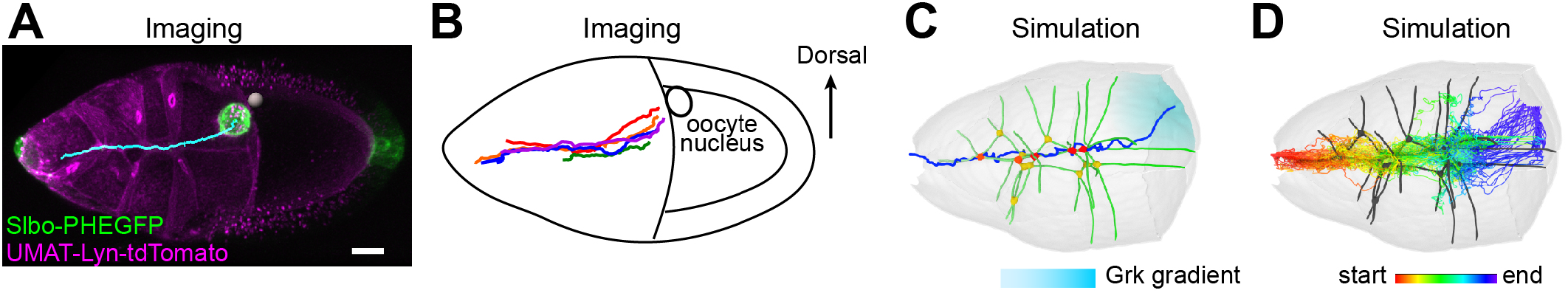
Integrating and prioritizing topographic and chemoattractant cues. (A) Live imaging of a wild-type egg chamber, showing dorsal migration near the oocyte. The cyan line indicates the track taken by border cells (movie 9). White sphere indicates dorsal location of the oocyte nucleus. Scale bar, 20 μm. (B) Dorsoventral alignment of egg chambers according to the oocyte nucleus position, by rotation around the x-y plane, reveals consistent dorsal migration near the oocyte. Trajectories from each of 5 egg chambers are shown in different colors. (C) Representative simulation of a border cell migration path when the Grk gradient (cyan shading) is included (movie 10). Note that between the most posterior four nurse cells, >3-cell junctures (colored dots) are absent, weakening the geometry cue and allowing the chemical cue to guide cells through a suboptimal physical space. (D) 99 simulations with Grk gradient.

## Discussion

Here we measure and manipulate chemical, adhesive, and topographical cues and elucidate their relative contributions to selection of one migration path amongst many. RTK signaling normally attracts border cells posteriorly toward the highest ligand concentration. We previously showed that E-cadherin amplifies small differences in chemoattractant concentration between the front and back of the cluster to ensure robust posterior migration (17). Here we show that the key function of nurse-cell E-cadherin is to provide traction. However, differential adhesion does not provide directional information to steer the cells.

For medial path selection, the organization of the nurse cells is an instructive cue. At the junctures where multiple nurse cells meet, they do not quite touch due to geometry, leaving tiny openings where protrusions need not break as many adhesion bonds between nurse cells. Thus, the concentration of multiple-cell junctures near the egg-chamber center provides an energetically favorable medial path. When the chemoattractant concentration is high enough, for example when the cells near the oocyte or in the presence of ectopic PVF1, the chemical cue can dominate, allowing cells to move through suboptimal physical space. Similarly, when E-cadherin-mediated traction is unavailable on nurse cells, border cells migrate on follicle cells, choosing grooves where multiple cells meet. This work thus elucidates how border cells integrate and prioritize chemical, adhesive, and physical features of the *in vivo* microenvironment to choose a path.

## Materials and Methods

### Drosophila genetics

Fly strains used in this study are listed in Table S1. Detailed fly genotypes in each experiment are listed in Table S2-3.

### CRISPR knockin of HA tag in mature chemoattractant ligand peptide

Sequences of PVF1, Krn, and Spi were aligned to determine HA insertion sites in the predicted mature peptide after cleavage (Tables S4). gRNA was designed (http://targetfinder.flycrispr.neuro.brown.edu/) after sequencing the host genomic DNA and cloned into the pU6-BbsI-chiRNA vector (Tables S5-6). HA tag insertion was designed according to scarless editing (https://flycrispr.org/scarless-gene-editing/) by cloning ~1kb left and right recombination arm into the pHD-2xHA-ScarlessDsRed vector. Primers used were listed in Table S7. CRISPR injection was performed by BestGene into fly strains expressing Cas-9 in the germ-line (Table S1 and S8). Individual F1 was crossed with PBac to remove scarless, and screened by eye color. Genomic DNA was extracted and sequenced to verify correct insertion. Egg chambers were stained live or fixed to detect HA signals (Table S8).

### Egg chamber dissection and staining

Adult female ovaries were dissected in Schneider’s Drosophila medium (Thermo Fisher Scientific, Waltham, MA; 21720) with 20% fetal bovine serum. Ovarioles containing egg chambers of the desired stages were pulled out of the muscle sheath with #55 forceps.

For fixed sample staining, ovarioles were then fixed for 20 min in 4% paraformaldehyde at 4 °C. For lectin (Lectin PNA Alexa 647; Thermo Fisher Scientific; L32460) staining, ovarioles were incubate in dissection medium containing 10 μg/ml lectin for 3 min before fixation. After fixation, ovarioles were washed with PBS/0.4% Triton X-100 (PBST), and then incubated with primary antibodies overnight at 4 °C. The following day, ovarioles were washed with PBST before incubation in secondary antibody for 1.5 h. After removal of secondary antibodies, samples were stained with Hoechst for 20 min. Samples were stored in VECTASHIELD (Vector Laboratories, Burlingame, CA) at 4 °C.

For HA-Keren live staining, egg chambers older than stage 10 were removed, and ovarioles incubated in live imaging medium containing insulin and anti-HA DyLight 550 (Thermo Fisher Scientific; 26183-D550) and Alexa-Fluor 647 (to mark non-specific trapping of antibodies in extracellular spaces) for 30 min. They were quickly rinsed two times with dissection medium and immediately mounted for live imaging. For dextran live labeling of extracellular space, dextran is membrane impermeant and readily diffuses between cells, but is endocytosed over time. Therefore we incubated living egg chambers in fluorescently labeled dextran (100 μg/ml; Dextran Alexa 647 10,000 MW; Thermo Fisher Scientific; D22914) and imaged them immediately, before endocytosis could occur.

Additional antibodies/dyes used in this study: rat anti-Ecad from Developmental Studies Hybridoma Bank (DSHB, Iowa City, IA; DCAD2; 1:50), chicken anti-GFP (Abcam, Cambridge, UK; 13970; 1:2000), Phalloidin for F-actin (Sigma-Aldrich; 65906; 1: 1000).

### Confocal imaging and visualization

For confocal z-stack imaging, in order to preserve the 3D structure and keep the sample stable during imaging, we adjusted the mounting and z-stack imaging method. First, for mounting, we used an 18 mm x 18 mm coverglass on the top and a 22 mm x 40 mm coverglass on the bottom for the inverted microscope to minimize sample movement by the touching objective during z-stacks. Four ~0.5 mm X 0.5 mm No.1 coverglass debris were used as bridges on four corners to avoid crushing of the egg chambers. 48.6 ul VECTASHIELD were used to allow 150 μl gap between the two coverglases so that the egg chambers were not compressed. Second, for imaging, we used a 40 × 1.1 N.A. 0.62mm long working distance water objective on a Zeiss LSM780 confocal microscope. Laser power corrections were applied by increasing the laser power as the objective scans from the near end of the sample to the far end of the sample.

Confocal z-stack images were visualized in Imaris (Bitplane, South Windsor, CT), Representative images were exported from Imaris using either 3D view or slice view. In Fig. 2H and I, Fig. 3I, and fig. S9A, nurse cell nuclei signals were masked to aid visualization of the membrane signals. In Fig. 3F and H, Fig. 5A, and fig. S14B, dextran signals outside of the egg chamber were masked to aid visualization of signals inside. Exported images were rotated and cropped in Photoshop (Adobe, San Jose, CA). Single channel images were converted from a black background to a white background using Invert LUT function in FIJI. Rainbow display of migration was generated using a subset of live image MIPs colored in FIJI using a custom LUT.

### Quantification in 2D, 3D and 4D

Keren Quantification: Antibodies can become stuck in extracellular spaces, as well as endocytosed by the border cell cluster. In order to correct for these non-specific fluorescent signals we incubated the egg chambers in both HA-Keren and Alexa-Fluor 647. The non-specific endocytosed Alexa-Fluor 647 signals were subtracted from HA-Keren channel. Final analysis was performed on the subtracted image of HA-Keren - Alexa-Fluor 647. The intensity of Alexa-Fluor 647 channel was attenuated by a factor calculated by comparing relative signal strength between HA-Keren and Alexa-Fluor 647, so that the subtracted HA-Keren retained the basal level intensity. In order to quantify the anterior-posterior signal, we drew a line with a width of 30 pixels in FIJI along the anterior-posterior local nurse cell membrane path. In order to quantify the medial-lateral signal we started from the lateral edge of the nurse cell membrane path and measured towards the medial center using the same method described above. We avoided quantifying the membranes that are curved, fuzzy, or are focused on ring canals’ focal planes. Raw distance was converted to the relative 0%-100% position along the path. Each data point of medial-lateral signal was normalized to the average within each egg chamber. Slope of 0%-100% medial-lateral normalized intensity was calculated across n=6 of each genotype, P value was based on t.test on the slopes. Each data point of anterior-posterior signal was normalized to the average of data points located between 40%-60% of the anterior-posterior migration path, at which every egg chamber has data points. Slope of 0%-70% anterior-posterior normalized intensity was calculated on three shgRNAi egg chambers and four control egg chambers that have all the data points on 0%-70%, and P value was based on t.test on the slopes. Data points from six to seven egg chambers are shown for each genotype. A fitted line represents the trendline of normalized Keren distribution along the anterior-posterior or medial-lateral axis. The standard deviation of the best fit trendline was represented by color shades in the graph.

For quantification of posterior and medial migration index in 3D, we used Imaris MeasurementPro in the following steps: 1) use an Oblique Slicer across the center of the nurse cell-ooctye border to set a plane parallel to the posterior follicle cell ring, set point B at the center of the nurse cell-oocyte border (fig. S1A); set point A at the anterior end of the egg chamber (Line AB measures and length of the anterior-posterior distance in nurse cells where border cells need to migrate); 2) move the previous Oblique Slicer going across the center of the border cell cluster, set point C at the intersection of the plane and line AB (Line AC measures how much border cell migrates from anterior); 3) on the Oblique Slicer plane that goes across C, set point D at the center of the border cell cluster, set point E and F at the edge of the egg chamber such that line EF goes through both the center of the cross sectional plane and the center of the border cell cluster (Line EF measures diameter of the egg chamber where the border cell is at). Posterior migration index = AC/AB * 100%; Medial migration index = min (DE, DF)/(EF/2) * 100%.

Posterior migration was quantified in stage 10 egg chambers. Note that in nurse cell Ecad knockdown, centripetal cells show migration defects so they are no longer the criteria for stage 9/10. Instead, use egg chamber size and nurse cell/oocyte ratio to distinguish stage 9/10. Medial migration was quantified in stage 9 egg chambers in which border cells had migrated 25-75% of the way to the oocyte border, where the abundance of alternative paths is greatest, except for Fig. 7 where we analyzed 75-100%. The preference for multiple nurse cell junctures was also evident at the initiation of migration (Fig. 5F to I).

For quantification of nurse cell membrane deflection in 3D, we used Imaris FilamentTracer in the following steps: 1) egg chamber drift in time was corrected by locating a few immobile anterior stretched cells, stalk cells, and posterior columna cells using the Spots function in Imaris Track based on Ubi-HisRFP; 2) The same 3-cell juncture was tracked each time frame using the AutoDepth method in Filaments based on the UMAT-Lyn-tdTomato signal, and were trimmed to ~8 μm length; 3) The Dendrite Orientation Angle was exported and used for calculate membrane deflection angle.

For quantification of >=3-cell juncture density in 3D, we used Imaris MeasurementPro in the following steps: 1) rotate the egg chamber in 3D so that it is not tilted in Z and the anterior is towards the left; 2) measure the anterior tip and the oocyte border position X, and then calculate 20, 30, 40, 50, 60, 70, 80% locations in X; 3) set a YZ Ortho Slicer in the each of these planes, add Measurement Points at the center of each YZ plane, as well as all the 3-cell junctures (Fig. 3B, blue dots); 4) in 3D view, trace all 3-cell juncture lines using Filament, and place Spots at the junctures of those filaments as >3 cell junctures (Fig. 3A, magenta dots); 5) export statistics for position XYZ, then calculate the distance to the center in the YZ plane; 6) bin the data in 5 μm radius intervals, and plot the heat map.

For quantification of dextran-labeled free space in 3-cell junctures (fig. S12E), we measured their diameter in 2D. An oblique slicer was used to find the 3-cell junctures that were close to the objective and almost parallel to the imaging plane to minimize the distortion from out-of-focus light. The line width of the 3-cell junctional space was measured from the slice using FIJI. Dextran-labeled free space in >3-cell junctures were measured in 3D. We used Imaris MeasurementPro Surface function and exported Volume statistics.

For quantification of 2-cell surface contact area in 3D, we used Imaris Cell function in the following steps: 1) rotate the egg chamber in 3D so that it is not tilted in Z and perform attenuation correction to adjust membrane signal intensity in Z; 2) Locate border cells, nurse cells, and oocyte nuclei using Spots function, add an additional Spot outside of the egg chamber, mask the nuclei stain channel with the Spots; 3) segment border cells, nurse cells, and oocyte using the Cell function by two channels, the masked nuclei channel and the membrane channel; 4) export Cells to Surfaces, and split nurse cell membrane surfaces and label them according to the identify in fig. S10A, S13A and S16C; 5) run Surface surface contact Macro in ImarisXT to segment different surface contact; 6) group them according to their location, and export statistics for the Area.

For quantification of the protrusion frequency in 4D, we did in the following steps: 1) manually set Spots at the tip of the protrusion by rotating the border cells in 3D in each time frame; 2) use an Oblique Slicer going perpendicular to the protrusion to determine if the protrusion is located at 2-cell interface or 3-cell juncture; 3) record the type of protrusion across the time frames.

For quantification of border cell cluster migration speed in 4D, we followed a published protocol (24): 1) egg chamber drift in time was corrected as described above; 2) border cell cluster was tracked using the Surfaces function in Imaris Track based on slbo-4XPHEGFP; 3) instantaneous speed was exported and plotted.

### Light sheet imaging and data processing

For light sheet imaging, younger and older egg chambers in the same ovariole of the targeted early stage 9 egg chamber were removed carefully with sharp forceps so they do not block the light path. The desired egg chambers were mounted in 0.5% low melting point agarose with fluorescent beads (Thermo Fisher Scientific; T7280) in 0.68 mm diameter capillary (Zeiss; 701902). Egg chambers that are oriented vertically in the capillary were chosen for imaging to minimize light blocking effect from the opaque oocyte. Images were taken from 8 angles 45 degrees apart on a Zeiss Lightsheet Z.1.

Images were registered, fused and deconvoluted using the Multiview-Reconstruction FIJI plugin (25). The fused and deconvolved egg chambers were then segmented into different germ cells using the Imaris Cell software and exported as surfaces. We then imported the surfaces into Meshlab to clean up the mesh as well as generate a PLY file that can be analyzed using Tissue Cartography (26). This was done by the following Meshlab commands: 1) importing the cell surface; 2) poisson disc sampling with base mesh subsampling; 3) computing normals for point sets; 4) surface reconstruction: poisson. The reconstructed surface is then exported as a PLY file. The PLY file was then analyzed using Tissue Cartography, details provided in the reference above. In short, the mesh is used to define the center of a region of interest. We then look at a sum of all of the pixels within a 1.5 um distance from the mesh. This signal is then “pulled-back” into a 2D image that can be analyzed using standard processing techniques. These 2D images were manually segmented and quantified in FIJI. Surfaces containing ring canals were not analyzed, due to both higher signal in the ring canal structure, and lower signal within the ring.

### FRAP imaging and data processing

Fluorescence recovery after photobleaching (FRAP) was done using a homozygous viable GFP trap line in the shotgun (Ecad) locus. Egg chambers were dissected in Schneiders media supplemented with 0.4 mg/mL insulin (Sigma) and 1X antimycotic/antibiotic (Gibco) before mounting on lumox dishes. FRAP performed using an inverted Zeiss LSM800 confocal microscope fitted with a Plan-Apochromat 40X, 1.2 NA multi-immersion objective with the collar set for water immersion. The Zeiss Zen software bleaching module was used to bleach a 3×5 μm rectangle of nurse cell membrane for 10 iterations using a 10 mW 488 nm diode laser with 3.1 mW at the sample plane. Image capture settings were as follows: 512×512 pixels (79.86 μm2 in XY) with a 316 msec scan time (1.03 μsec pixel dwell) using a 3.93 Airy unit or 3.2 μm section thickness at a bit depth of 16. A total of 5 pre-bleach images were taken at a 2.5 sec interval followed by 3 min of post-bleach imaging at the same 2.5 sec imaging interval. Fluorescence intensity was measured for the bleached membrane region, a nonbleached control membrane region and a background region outside of the egg chamber. These values were then input into the easyFRAP web application (https://easyfrap.vmnet.upatras.gr) to calculate the T1/2 and mobile fraction of E-cadherin on side versus center nurse cell membranes. Graphs were then processed in Prism (GraphPad, La Jolla, CA).

### Statistics and reproducibility

All fly crosses were repeated at least twice and ovary dissections and staining were repeated at least three times. Data describe technical or biological replicates. Sample size was not predetermined by statistical methods but we used prior knowledge to estimate minimum sample size. Samples of the correct stage were included and no data point that fits the analysis criteria was excluded. The experiments were not randomized. Investigators were not blinded.

Standard statistical tests were performed using Prism. Sample sizes were appropriately large with appropriate distributions. Mann–Whitney nonparametric test (two-tailed) was used for comparing two groups with different variance. Kruskal–Wallis nonparametric test, followed by Dunn’s multiple comparisons test was used when the variance is significantly different among multiple groups. Paired t-test (two-tailed) was used for comparing the two measurements from the same sample.

### Data and code availability

The authors declare that all data and code supporting the findings of this study are available within the article and its supplementary information files or from the corresponding author upon reasonable request.

## Acknowledgments

We thank Benjamin Cheng, Shwena Dhar, Yijing Li, Marc Anthony Pastor and Dr. Rui Wu for technical assistance. Funding: This work was supported by NIH grant GM46425 to D.J.M, NSF grant PHY-1707637 to W.J.R. and ACS grant PF-17-024-01-CSM to J.P.C. We thank the Developmental Studies Hybridoma Bank for providing antibodies and Drs. Hsin-Ho Sung, Huynh Jean Rene, the Bloomington Drosophila Stock Center, and the Vienna Drosophila Resource Center for providing fly stocks. We acknowledge the use of the NRI-MCDB Microscopy Facility and the Imaris computer workstation supported by the Office of The Director, National Institutes of Health of the NIH under Award # S10OD010610.

## Author Contributions and Notes

Experiments were designed by W.D., X.G., J.P.C, J.A.M. and D.J.M. Experiments were carried out by W.D., X.G., J.P.C, J.A.M., and H. B. Modeling was carried out by Y.C., W.J.R. and N.G. Graphical illustrations and animations were produced by B.J.M. S. S. assisted with light sheet imaging and data analysis. The manuscript was prepared by W.D., X.G.,J.A.M., J.P.C., Y.C., W.J.R., N.G. and D.J.M.

The authors declare no conflict of interest.

All data are available in the manuscript or the supplementary materials. Materials available upon request.

## Supplementary Materials

Supplementary Text Fig. S1 – S16 Tables S1 – S11

Multimedia Files: Movies 1-10 References (24–29)

## List of Supplementary Materials

### Supplementary Text

#### ST1. The free space in 2D and 3D

To determine the possible free space in a N-cell juncture, we assume that the equilibrium configuration is determined by a balance between the adhesion energy between nurse cells (parameterized by A, per unit length in 2D, and per unit area in 3D) and the membrane/cortex bending energy (parameterized by bending modulus B, per unit length in 2D and per unit area in 3D). First, we consider the 2D case shown in fig. S12 A and B for *N* = 3 and *N* = 4. The geometry of the junctures is characterized by circular arcs of membrane with radius *R_f_* that join smoothly, i.e. without creating a cusp. The system’s energy reaches a minimum when *δ*(*W_adh_* + *W_bend_*)/δ*R* = 0. The bending energy for each arc is computed by integrating over the angle of polygon (*N*-2)*π/N* (for the examples shown in fig. S12A and B this angle is π/3 and π/2, respectively). Computing the energy change for each arc, we arrive at δ*W_bend_* = −*BN*θ(*N*)δ*R*/*R*^2^ with θ(*N*) = (*N* – 2)π/*N*. Due to the change in *R* the total length of cell-cell contact changes and the resulting total change in adhesion energy is δ*W_adh_* = *NA*δ*L* = *N*(*A* cot[θ(*N*)/2]δ*R*). Setting δ(*W_adh_* + *W_bend_*) = 0 we find for the equilibrium radius

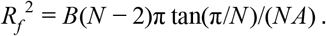

The area of the free space is then the area of the N-polygon connecting the circle center minus the area of the circular sectors. The area of the polygon is 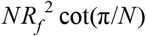 while the area of sectors is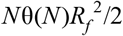. Thus, the total free space is given by

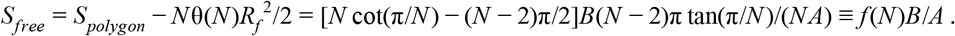

In fig. S12C we show that *f*(*N*) = *S_free_*/(*B*/*A*) is a monotonically increasing function of N. This indicates that the available space becomes larger as the number of cells in the juncture increases. Note that *S_free_* ~ *B/A* which predicts that, as adhesion decreases, the free space in cell junctures increases, which is consistent with the results of the E-cadherin knock-down experiments (Fig. 3H, Fig. 5A, and fig. S12E).

Following the same argument, the above calculation can be directly extended to 3D, where the nurse cell membranes, now considered to be spherical caps, join smoothly at the juncture. Unlike 2D, where any number of arcs can join to form a polygon, in 3D the stacking of spherical caps is more complicated. Specifically, it is only possible for N=4,6,8,12, and 20, such that the center of the spheres form a polyhedron known as a Platonic solid. The N=4 example, resulting in a tetrahedron, is shown in fig. S12D. We again have the adhesion-bending δ(*W_adh_* + *W_bend_*)/δ*R* = 0, where close at δ*W_adh_* = *ANR*δ*R* cot[θ(*N*)/2] and δ*W_bend_* =−*BN*ψ(*N*)δ*R*/*R*^3^. Here, θ(*N*) and ψ(*N*) is the dihedral angle and the solid angle of the face of the polyhedron, respectively. The equilibrium radius is thus

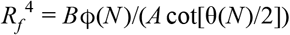

The free space volume is the volume of the Platonic solid minus the volume of the spherical caps,

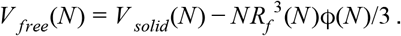

For N=5, a simple configuration is mirror imaging a tetrahedron (N=4), which gives *V_free_*(5) = 2*V_free_*(4). For N=7, there is no simple stacking. The ratio of the free space volume between N=4,5,6, and 8 in 3D is 1 : 2 : 6.6 : 11.6, which increases as N increases. This is close to the experimental ratio, which was found to be 1 : 3.8 : 9.5 : 15.5 (Fig. 3G, the middle point is arbitrary and is scaled to match the experimental data). As a comparison, the ratio in 2D is N=4,5,6,8 is 1 : 2.3 : 3.8 : 7.4.

#### ST2. Energy costs of protrusions

When border cell protrusions extend into a two-nurse-cell juncture, they have to “unzip” the adhesion bonds between nurse cells. Additionally, there is an energy cost due to the need to bend nurse cell membranes (Fig. 3E, orange box). The adhesion penalty is approximately *W_adh,2_* = 2π*rA* ≈ *AP*, where *r* is the average radius of the border cell protrusion and *P* is its perimeter. The bending penalty is *W*_*bend*2,_ ≈ 2π*B*/*r*. When border cells protrude into *N* ≥ 3 cell junctures, the protrusions break fewer adhesion bonds due to the pre-existing space, so the adhesion penalty is smaller than in two cell junctures. Furthermore, in contrast to the two-cell juncture, squeezing into *N* ≥ 3 should *release* bending energy, by reducing the nurse cell curvature from 1/*R_f_* to 1/*R_p_*, where *R_f_* and *R_p_* represent the local radius of the nurse cell membrane before and after the protrusion moves in (Fig. 3E, cyan and magenta boxes). In general, *R_p_* > *R_f_*. The adhesion penalty is proportional to the nurse cell membrane that must be unzipped, given by *W_adh,N_* = *A*(*N* – 2)π(*R_p_* – *R_f_*). The difference of total bending energy is *W_bend,N_* = *B*(*N* – 2)π(1/*R_p_* – 1/*R*). In the simple case, we can model border cell protrusions in full contact with the nurse cells with a fixed perimeter of (*N* – 2)π*R_p_* = *P*. Note that, in this case, the adhesion difference due to nurse-border cell contacts is independent of *N*. Then, *W*_*adh*,2_ > *W_adh,N_* and *W*_*bend*,2_ > 0 > *W_bend,N_* for *N* ≥ 3. Thus, both bending and adhesion energies favor protrusion into *N* ≥ 3-nurse-cell junctures rather than into two-cell junctures. Furthermore, *W_adh,N_* = *A*(*N* – 2)π(*R_p_* – *R_f_*) = *AP* – *A*(*N* – 2)*R_f_* decreases as *N* increases. Values of *W_adh,N_* are listed in Table S9.

Furthermore, we estimate the value of *B/A* ≈ 1 μ*m*^2^ by fitting the experimental data in Fig. 3G with the free space volume ratio in 3D. The relative contribution from adhesion and bending energy can be found from 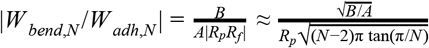. *R_p_* can be estimated from the size of the border cell protrusions 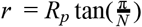. From experimental data, we can estimate *r* ≈2*μm* (Fig. 3J). Plugging the estimates we get *W_bend,N_*/*W_adh,N_*| ≪ 1 for *N* ≥ 3, which indicates that the bending energy is negligible compared to the adhesion energy.

#### ST3. The 3D dynamic model

To understand the dynamics of border cell cluster migration in response to physical and chemical cues, we developed a dynamic model that describes the trajectory of border cells within an experimentally determined three-dimensional topography of egg chambers. A complete physical model would describe border cells as deformable 3D objects that exert forces on each other and on deformable nurse cells, and that migrate through a complex 3D geometry guided by chemotactic and geometric cues. Such a model is currently not feasible. We, therefore, model the border cell cluster as a point particle, representing the center of mass, that moves in a 3D geometry based on experimental data, and that follows experimentally derived rules. This model allows us to probe the relative importance of topographic and chemotactic cues.

The mass center of the border cell cluster is modeled as a particle located at position 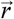 moving in an effective potential 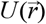 and subject to noise 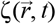. The motion can be described by an overdamped stochastic Langevin equation

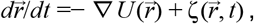

where 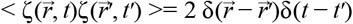. The effective potential incorporates the different guidance terms:

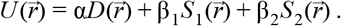

The topographic cue is described by the first term. It takes into account that the border cells have multiple protrusions which probe their surroundings. The border cell cluster is more likely to move into places where more free space is available. We model this by including the average energy cost in a sphere of cluster size

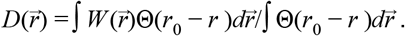

Here Θ(*r*_0_ – *r*) = 1 when *r*_0_ > *r*, otherwise Θ(*r*_0_ – *r*) = 0, *r*_0_ is the average radius of the border cell cluster, and 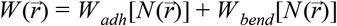, where *W_adh_* and *W_bend_* are the energy costs of protruding into a N-cell juncture at position 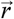 (computed in ST2). Note that 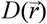 has a dominant weight to the outer surface of the cluster where protrusions are formed.

The chemotactic cues are described by the second and third term and take into account chemoattractant gradients along the anterior-posterior and dorsal-ventral axis. Along the anterior-posterior axis, we use, following (*27*), an exponential concentration profile: 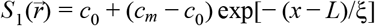, where ξ is the length scale of the exponential profile decay, and *L* is the total length of the egg chamber. Dorsal-ventral guidance is incorporated by a concentration gradient of the protein Gurken, which we represent as 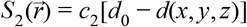, where 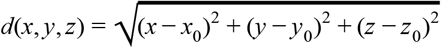 is the distance to the Gurken source. Earlier studies revealed that this guidance is only relevant near the posterior of the egg chamber and thus only exists in our model when *x* > 2*L*/3 (*5*). Furthermore, we assume that nurse cells represent static obstacles such that 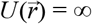 when 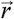 falls within a nurse cell. Finally, the strength of the geometric cue and chemotactic cue is determined by the parameters α, β_1_ and β_2_.

In our simulations, we first discretized space in an experimentally segmented egg chamber, resulting in a cubic grid of 1 μm. Each grid point is assigned with an identity which indicates the juncture type it belongs to (e.g., 2 or 3-cell juncture). Our simulations started by placing the particle (i.e., border cell cluster) at the anterior end of the egg chamber. We then incorporated a discrete version of the Langevin equation in which, for each time step, the particle can move to a neighboring cubic grid points with probability

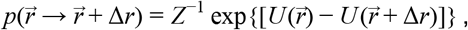

where 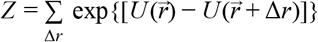 is the partition function, and the sum is over all nearest neighbors of position 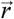. The potential can be parameterized by rescaling units such that *A* = 1 (notice that the bending energy is negligible, as shown in ST2). The simulation was terminated when the particle reached the oocyte or the total number of simulation steps exceeded 10^4^ (a typical simulation in wild type egg chamber takes ~ 10^3^ steps.). Parameters used in the model are given in Table S10. For different combinations of α and β, the mediolateral and anteroposterior indices are shown in Table S11.

Our model can also be applied to the case when E-cadherin is knocked down. In the E-cadherin knockdown egg chamber, the nurse-follicle cell junctures are 2.5 fold larger than the wild type nurse-follicle cell juncture (Fig. 5A, fig. S12E). This difference in juncture size indicates that the adhesion between nurse cell and follicle cells is smaller (see section above). In our simulation, we set *A* = 0.4 while keeping the other parameters the same.

### Supplementary Figures and Legends

**Fig. S1.**
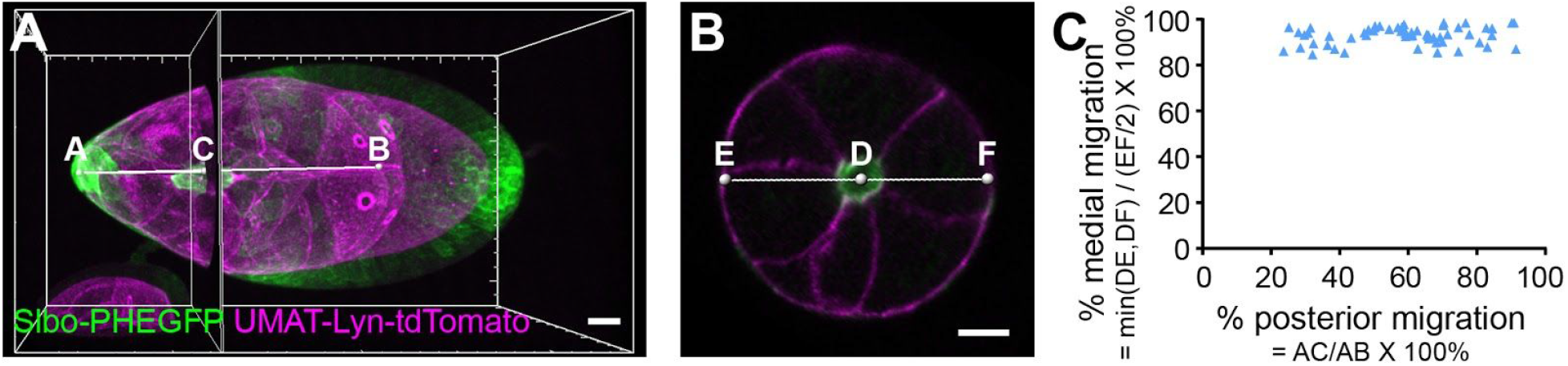
3D analysis reveals orthogonal anteroposterior and mediolateral pathway choices. (A) Lateral view showing the method for quantification of posterior migration. (B) Cross sectional view showing the method for quantification of medial migration. (C) Migration in control stage 9 egg chambers. (n = 59 egg chambers). Scale bars, 20 μm.

**Fig. S2.**
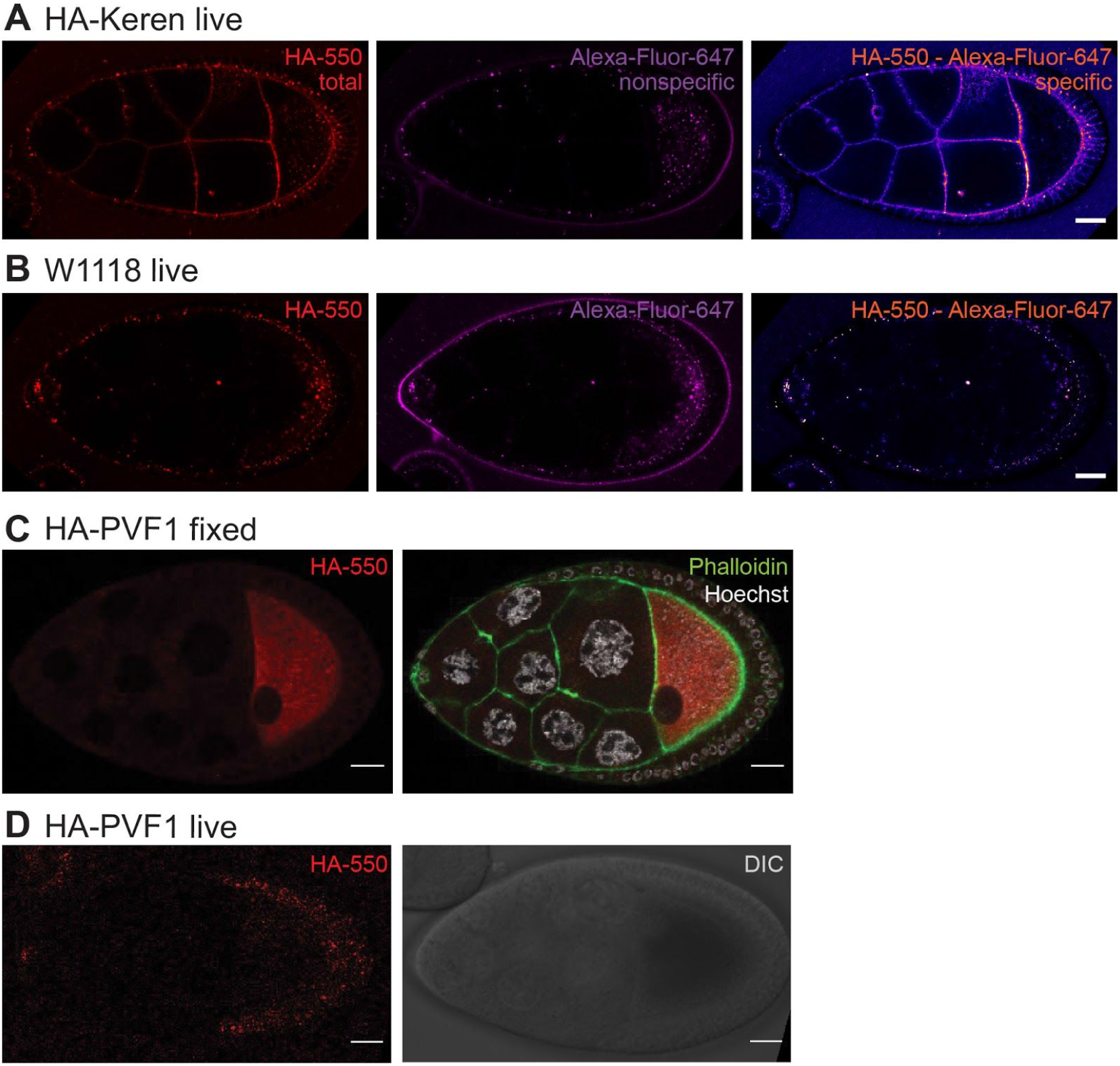
HA tagging chemoattractant staining. (A) Staining of a living stage 9 egg chamber from a homozygous CRISPR HA-Keren fly with anti-HA-550 and a non-specific Alexa-647 antibody. The 550 - 647 signal provides the specific HA-Krn signal. (B) Staining of a stage 9 egg chamber from w1118 (negative control) with the same antibodies as in A. (C) Staining of a fixed and permeabilized stage 9 egg chamber from a homozygous CRISPR HA-PVF1 fly with anti-HA-550 antibody alone and with Hoechst for DNA (white) and Phalloidin for F-actin (green). (D) Staining of a living stage 9 egg chamber from a homozygous CRISPR HA-PVF1 fly with anti-HA-550 showing that extracellular PVF1 is not detected, though the tagged protein is functional because homozygous flies are viable and fertile (Table. S8). DIC imaging of the same egg chamber is shown. Scale bars, 20 μm.

**Fig. S3.**
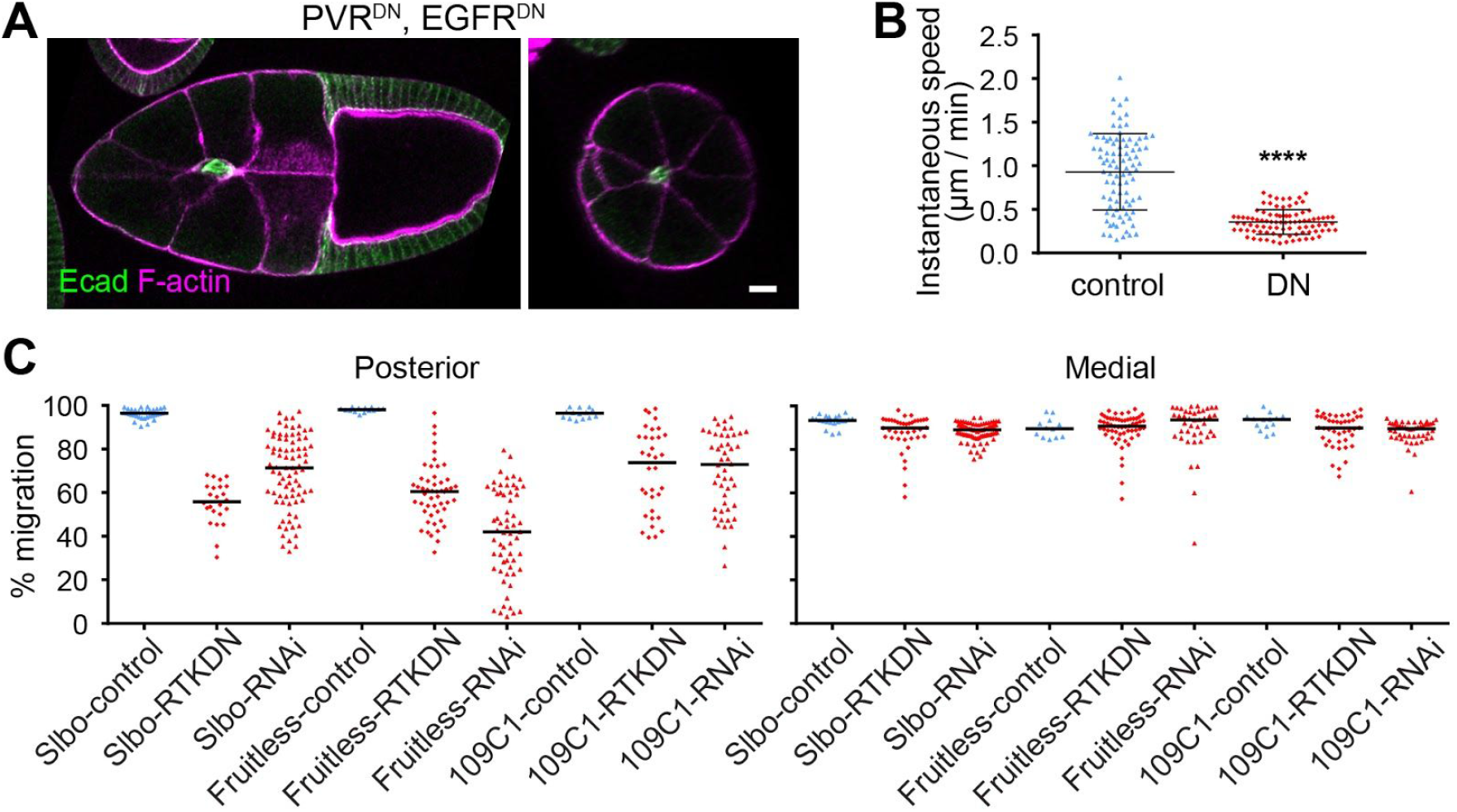
Migration defects caused by PVR and EGFR RNAi or dominant-negatives (DN) (A) Lateral and cross sectional view showing an example of posterior migration defect but normal mediolateral path selection in slbo-Gal4>UAS-PVR^DN^, EGFR^DN^. Scale bars, 20 μm. (B) Quantification of instantaneous migration speed during early stage 9, in 2 minute time intervals over the course of one hour in slbo-Gal4 control or slbo-Gal4>UAS-PVR^DN^, EGFR^DN^. Data from *n* = 90 time points (30 time points each from 3 egg chambers per group). ****, *P* < 0.0001 (Mann-Whitney test). Reduced migration speed shows the dominant-negative receptors were effective. (C) Quantification of migration index following expression of EGFRDN and PVRDN (RTKDN) with the following Gal4 lines: Fruitless-Gal4 expresses in border cells from stage 4, 109C1-Gal4 from stage 7, slbo-Gal4 from stage 8.

**Fig. S4.**
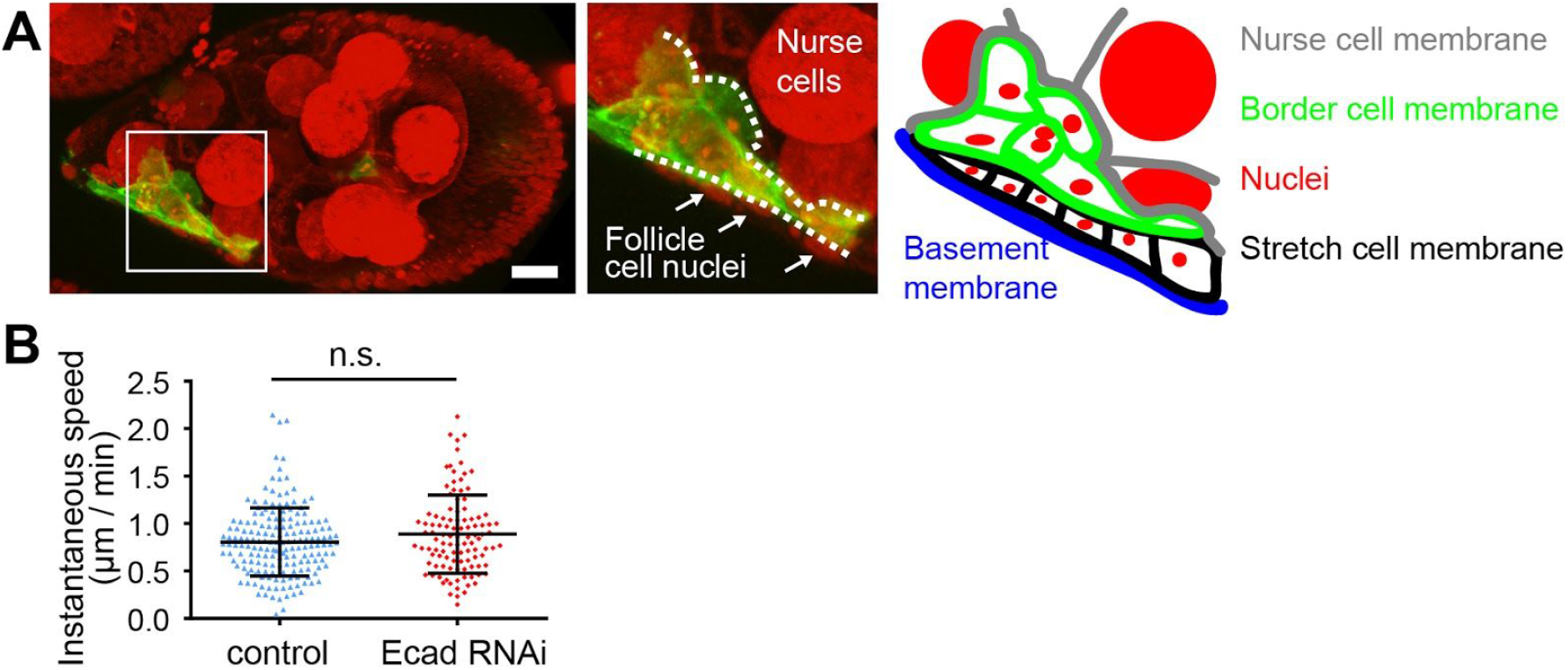
Effect of nurse cell Ecad knockdown on border cell migration. (A) In nurse cell Ecad knockdown, border cells (marked by Slbo-PHEGFP) migrate in between follicle cells (white arrows) and nurse cells. Scale bars, 20 μm. (B) Quantification of instantaneous migration speed every 4 minutes in control Matalpha-Gal4>UAS-wRNAi or Matalpha4-Gal4>UAS-Ecad RNAi. In contrast to RTK inhibition, migration speed was not initially reduced. Data from eight WT controls n = 178 time points; eight nurse cell Ecad RNAi egg chambers n = 107 time points.

**Fig. S5.**
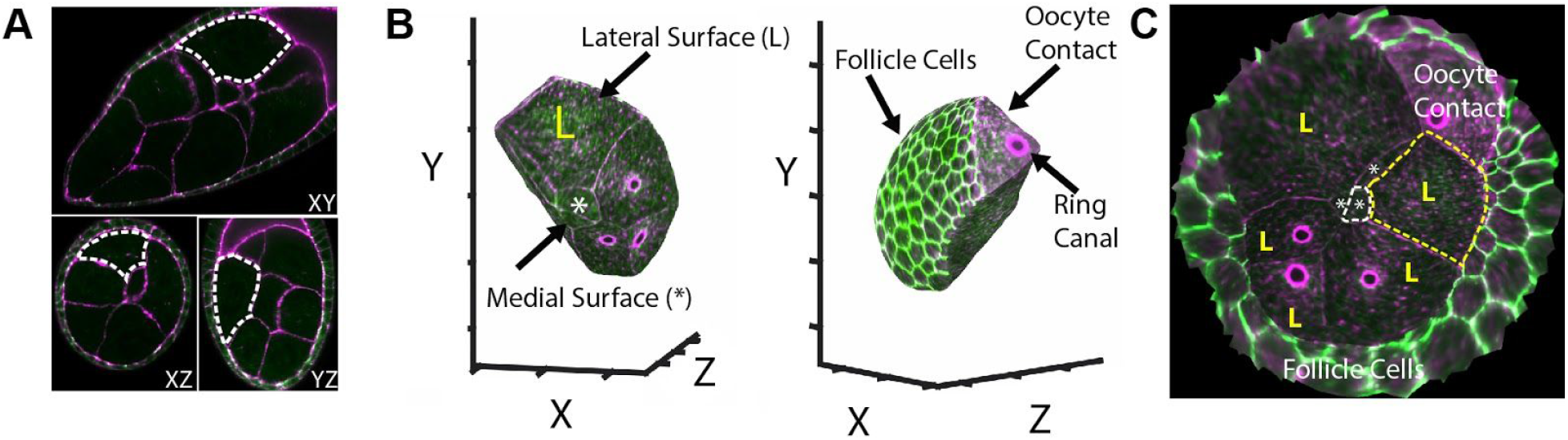
Isotropic light sheet imaging shows no differences between Ecad concentrations on medial vs. lateral membranes. (A) Isotropic light sheet imaging of a stage 9 egg chamber stained for F-actin with phalloidin (magenta), and anti-Ecad antibody (green). Dashed lines indicate the boundaries of a single nurse cell in the XY/XZ/YZ orthogonal views. (B) 3D reconstruction of the surface of the nurse cell outlined in (A). (C) “Pullback” of the surface of the 3D reconstruction in (B) created with the Images Surfaces Analysis Toolkit (*26*). The pullback represents the summed intensity projection of 1.5um thickness around the surface of the 3D object in (B). Medial surfaces are represented by (*) and lateral surfaces by (L).

**Fig. S6.**
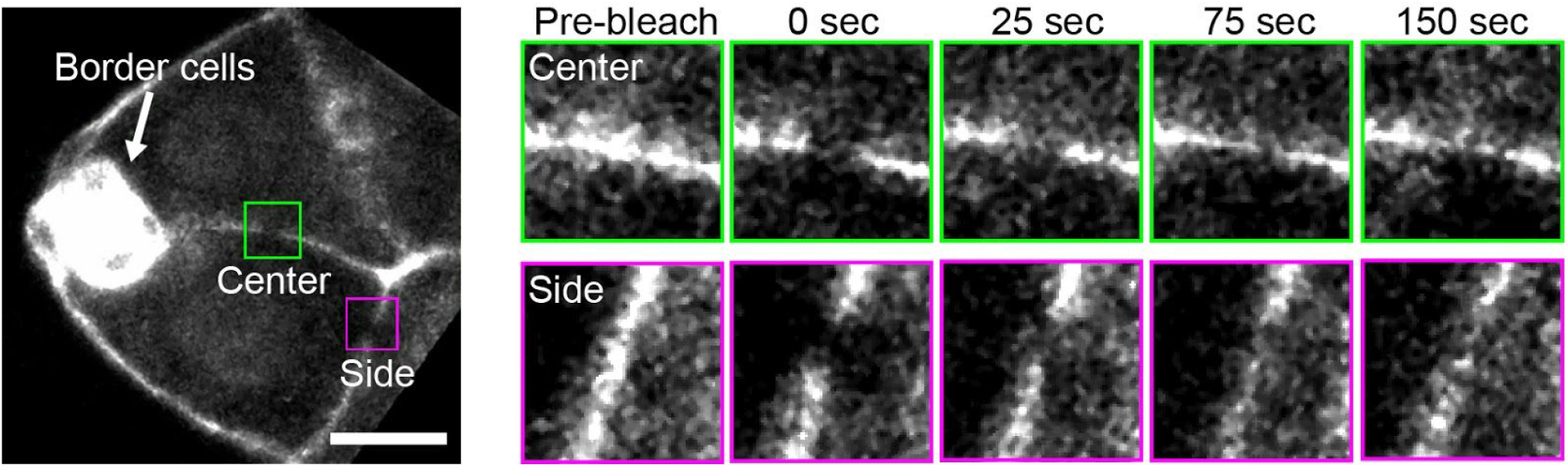
Fluorescence recovery after photobleaching (FRAP) shows no difference in Ecad stability on center and side paths. Stage 9 egg chamber with Ecad endogenously tagged with GFP. Insets show center vs. side membrane Ecad signal. Scale bar, 20 μm.

**Fig. S7.**
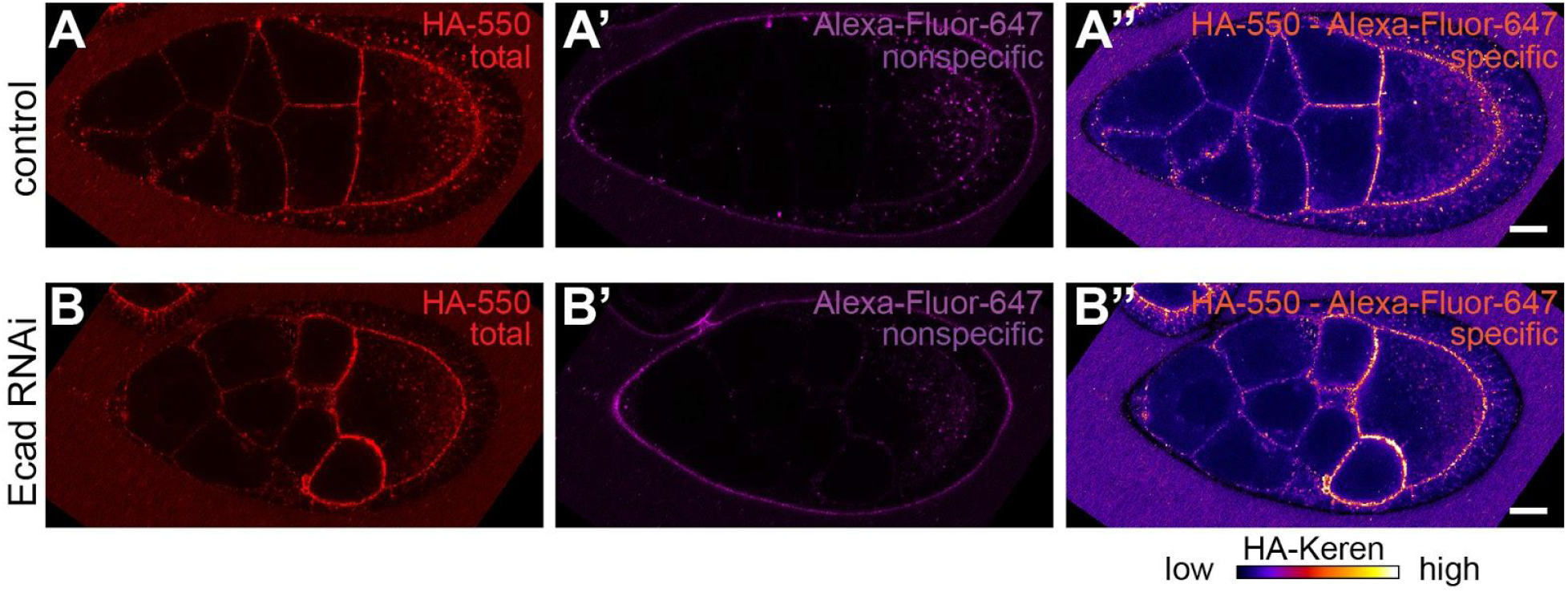
HA-Krn concentration in control and nurse cell Ecad RNAi expressing egg chambers. Ecad RNAi does not cause a redistribution of HA-Krn that would account for the mediolateral border cell guidance defect. (A) Confocal imaging of anti-HA-550 staining of a living stage 9 egg chamber from a heterozygous CRISPR HA-Keren fly (negative control) [genotype: MatalphaGal4/+;HA-Keren/+;UAS-wRNAi/+]. (A’) Labeling of the same egg chamber using a non-specific Alexa-647 antibody. (A’’) The specific pattern of HA-Krn was calculated by subtracting the non-specific 647 signal from the total 550 fluorescence. (B-B’’) Staining of a stage 9 egg chamber from MatalphaGal4/+;HA-Keren/+;UAS-EcadRNAi/+ using the same method as in A-A’’. Scale bars, 20 μm.

**Fig. S8.**
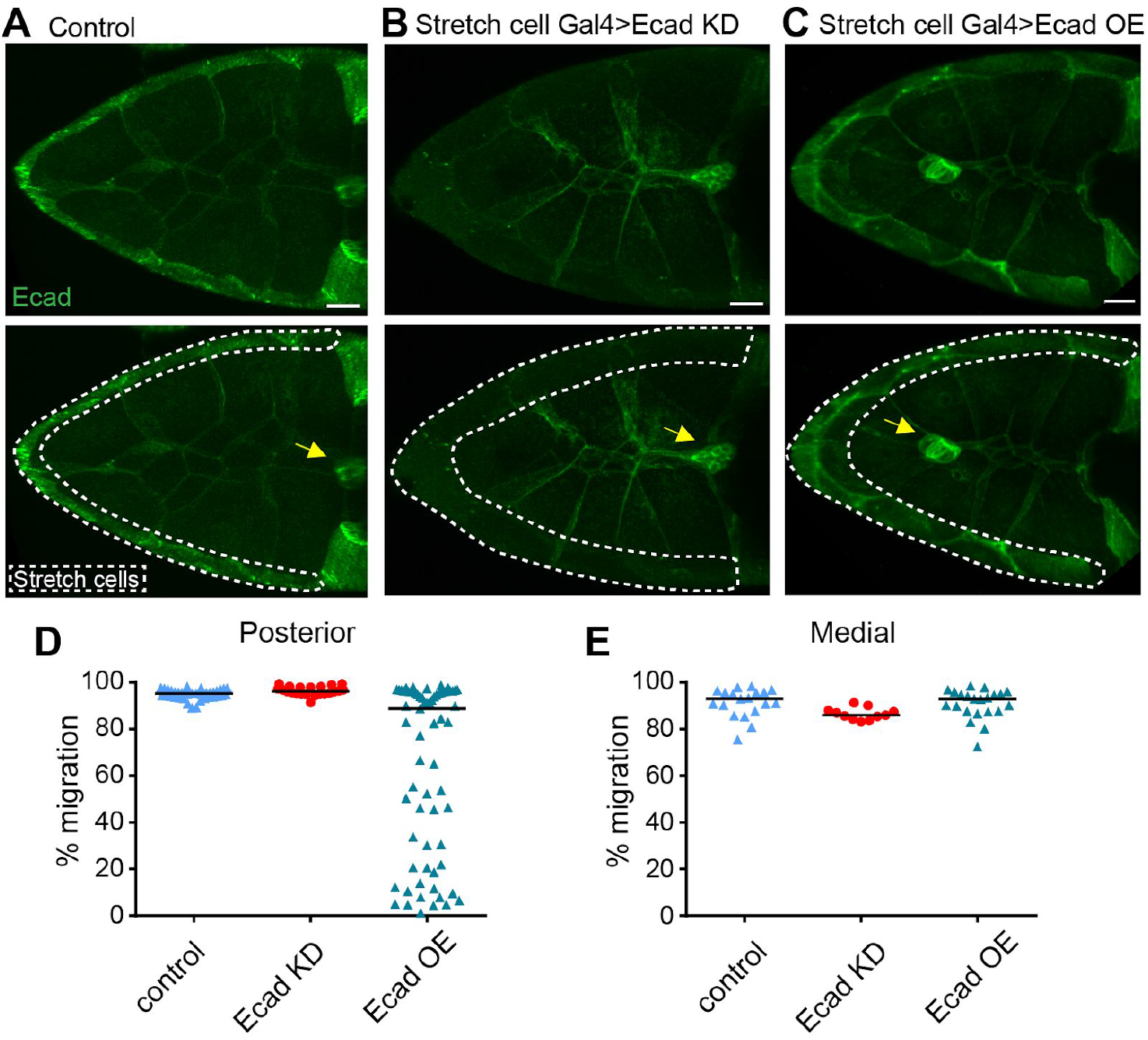
Stretch cell Ecad knockdown shows no medial guidance defect. Projections of 18 confocal sections (2um each) of anti-E-cadherin staining of early stage 10 egg chambers in the indicated genotypes. (A) Control, stretch cell Gal4>wRNAi, (B) Decreased Ecad level in stretch cell Gal4>Ecad RNAi. (C) Increased Ecad in stretch cell Gal4>Ecad. Scale bar, 20 μm. Dotted lines show stretch cell regions; Arrows indicate border cell clusters. Quantification of posterior (D) and medial (E) migration in egg chambers for each genotype. Scale bars, 20 μm.

**Fig. S9.**
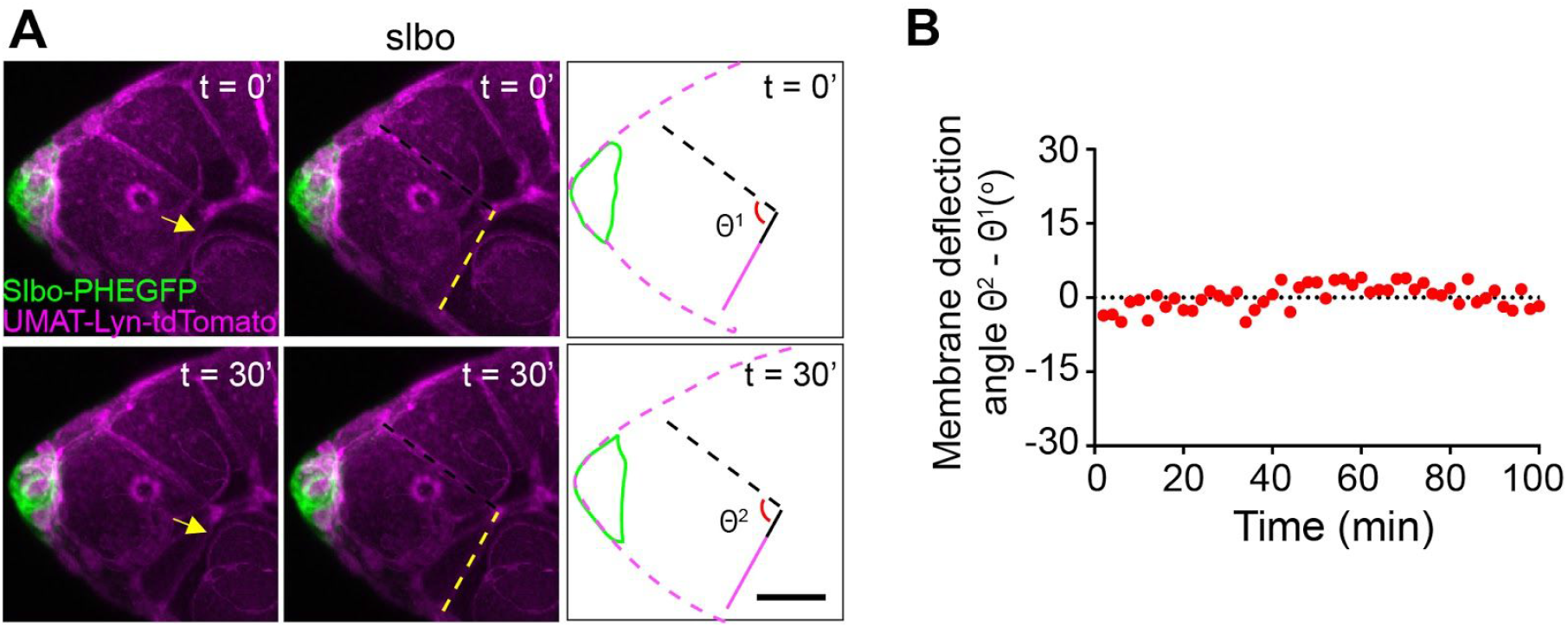
Nurse cell membrane deflections require border cell migration. (A) Still images from a movie of a *slbo* mutant egg chamber in which border cells do not migrate (*28*). No nurse cell juncture deflection occurs, showing that they are not random fluctuations; rather they are caused by border cells actively pulling. Scale bar, 20 μm. (B) Representative trace of normalized nurse cell membrane deflections in the *slbo* mutant.

**Fig. S10.**
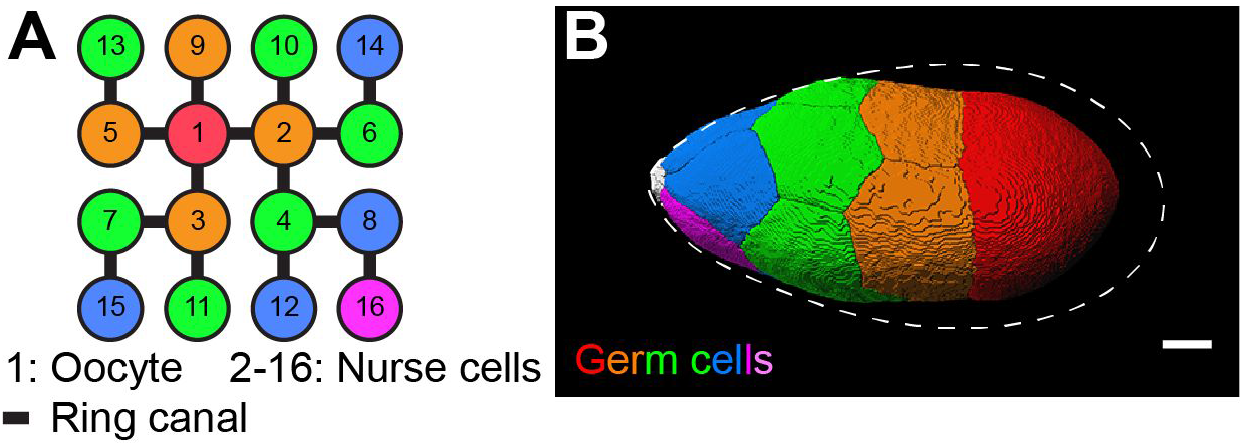
Nurse cell arrangements in control egg chambers. The 16 germ cells derive from a single germline precursor, which undergoes four rounds of cell division with incomplete cytokinesis. Residual, stabilized cleavage furrows called ring canals thus connect germline cells to one another in a regular pattern. (A) Schematic drawing of the germ cell identity (numbers in circles) based on their birth order and thus ring canal connections (lines) in control egg chambers. Colors indicate distance to the oocyte (#1). (B) 3D reconstruction of germ cell packing in stage 9 egg chambers. White, border cells at the anterior tip. Dashed line indicates the somatic follicle cell layer. Scale bar, 20 μm.

**Fig. S11.**
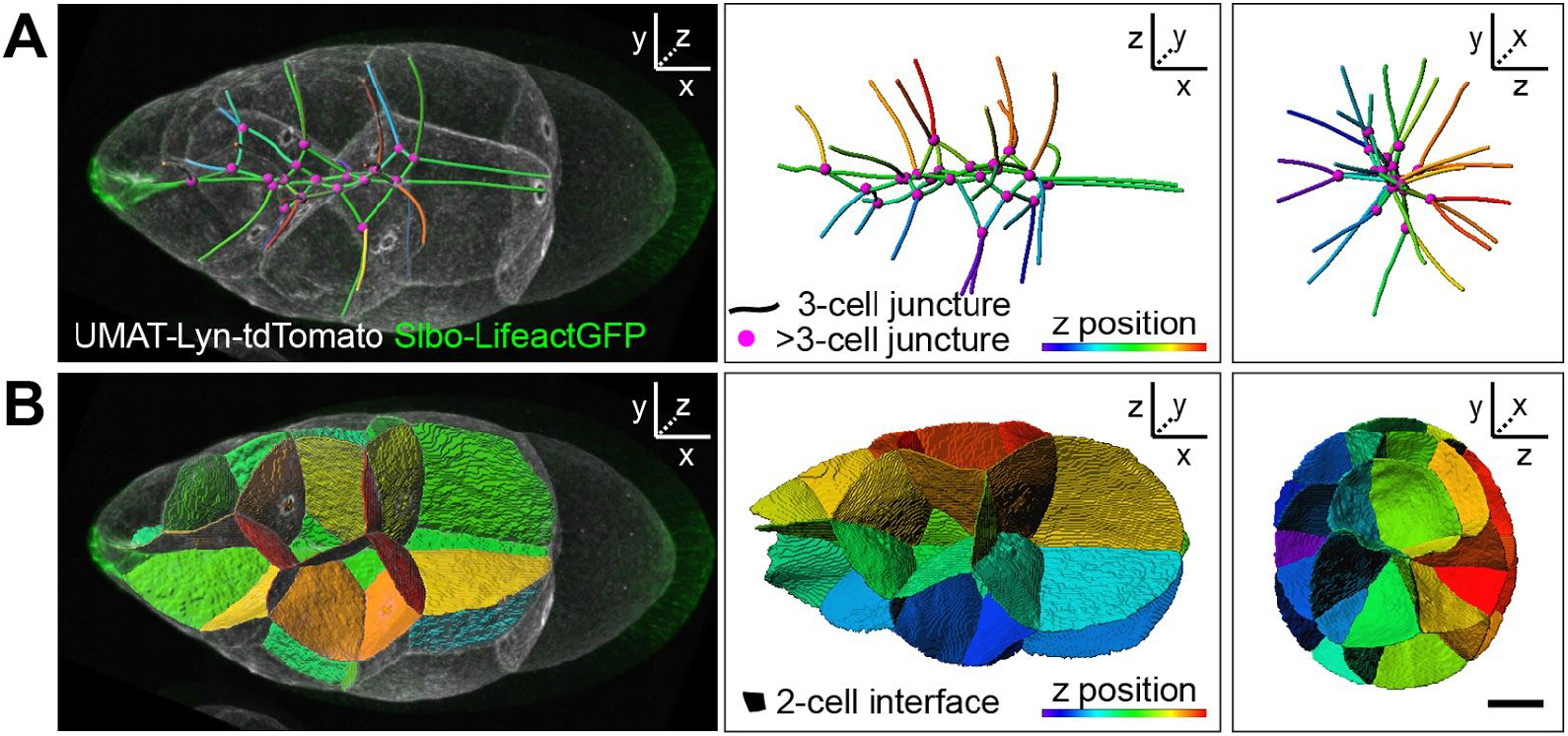
Three dimensional reconstructions of nurse cell contacts. A second example for Fig.3A and C. (A) 3-nurse cell junctures are represented as lines and color provides position in z. Magenta dots represent junctures of >3 nurse cells. x,y planes (left panels), x,z planes (middle), and y,z planes (right panels) are shown. (B) Two-cell-contacts are shown as surfaces. Scale bar, 20 μm.

**Fig. S12.**
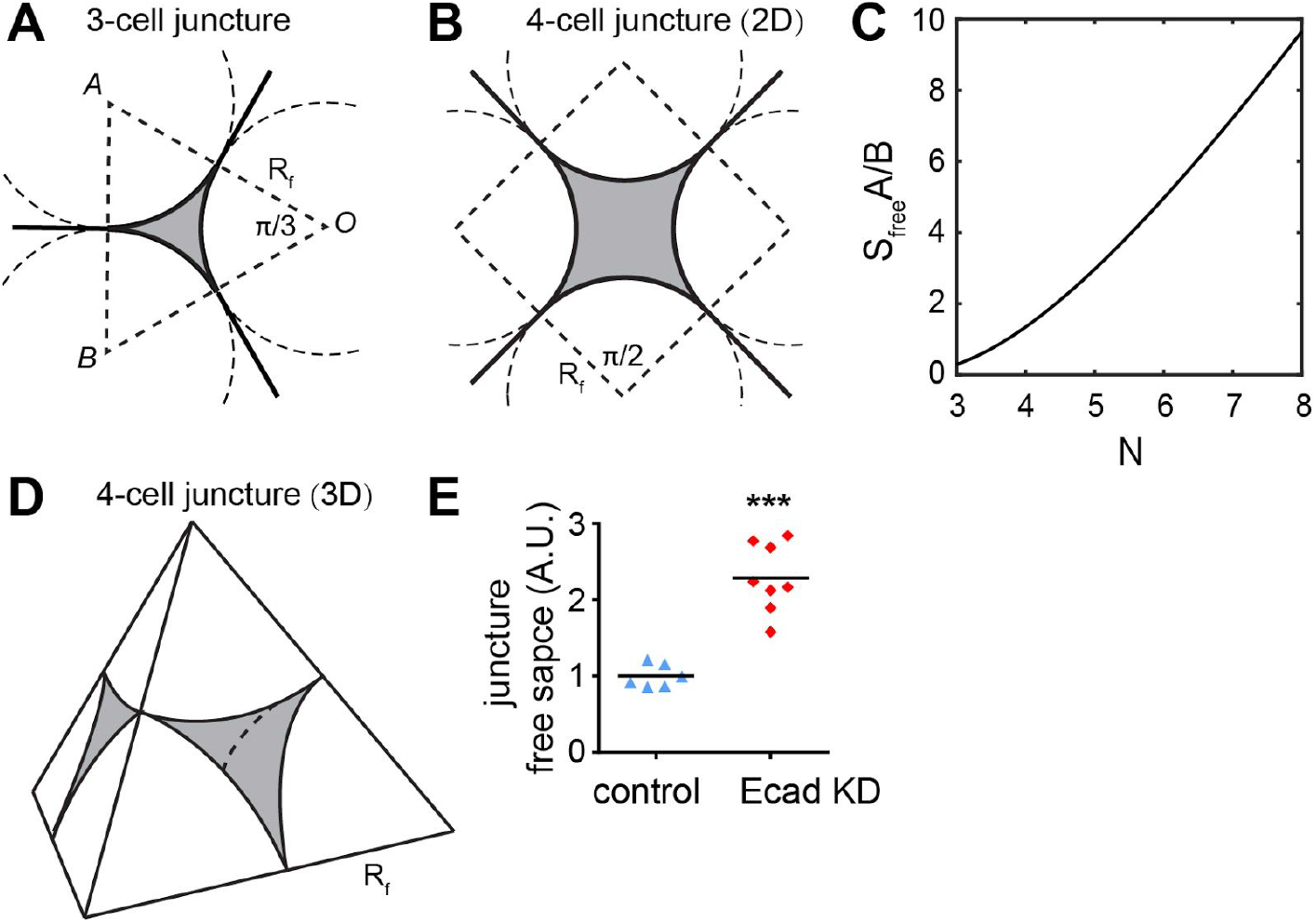
Geometry of nurse cell junctures and associated free space. (A) The local configuration of 3-cell junctures in a 2D cross section. Nurse cell membranes are represented by solid black lines and R_f_ is the local radius of the nurse cell. The free space between the nurse cells, shaded gray, is the area of the polygon (OAB), a triangle in this case, minus the area of the circular sector with centers at A, B and O, multiplied by the number of nurse cells in the juncture. (B) The configuration of a 4-cell juncture in 2D. In this case, the polygon is a square. (C) S_free_ is a function of the balance between adhesion (A) energy gained and the cost of bending (B) nurse cell membranes and increases with increasing N at the junctures. Bending energy proved negligible (see ST2) (D) The configuration of a 4-cell juncture in 3D. The nurse cell membrane is represented by spherical surfaces that join together. The centers of the spheres are the vertices of a tetrahedron, which has edges with length 2R_f_. The volume of the free space, again shaded gray, is the volume of the tetrahedron minus the volume of the spherical caps. (E) Quantification of 3-cell junctures in 2D. n = 6, 8 pairs of junctions from 3 control and 4 ECad KD egg chambers. Bars show mean. ***, *P* < 0.001 (Mann-Whitney test), A.U. arbitrary unit. Scale bar, 20 μm.

**Fig. S13.**
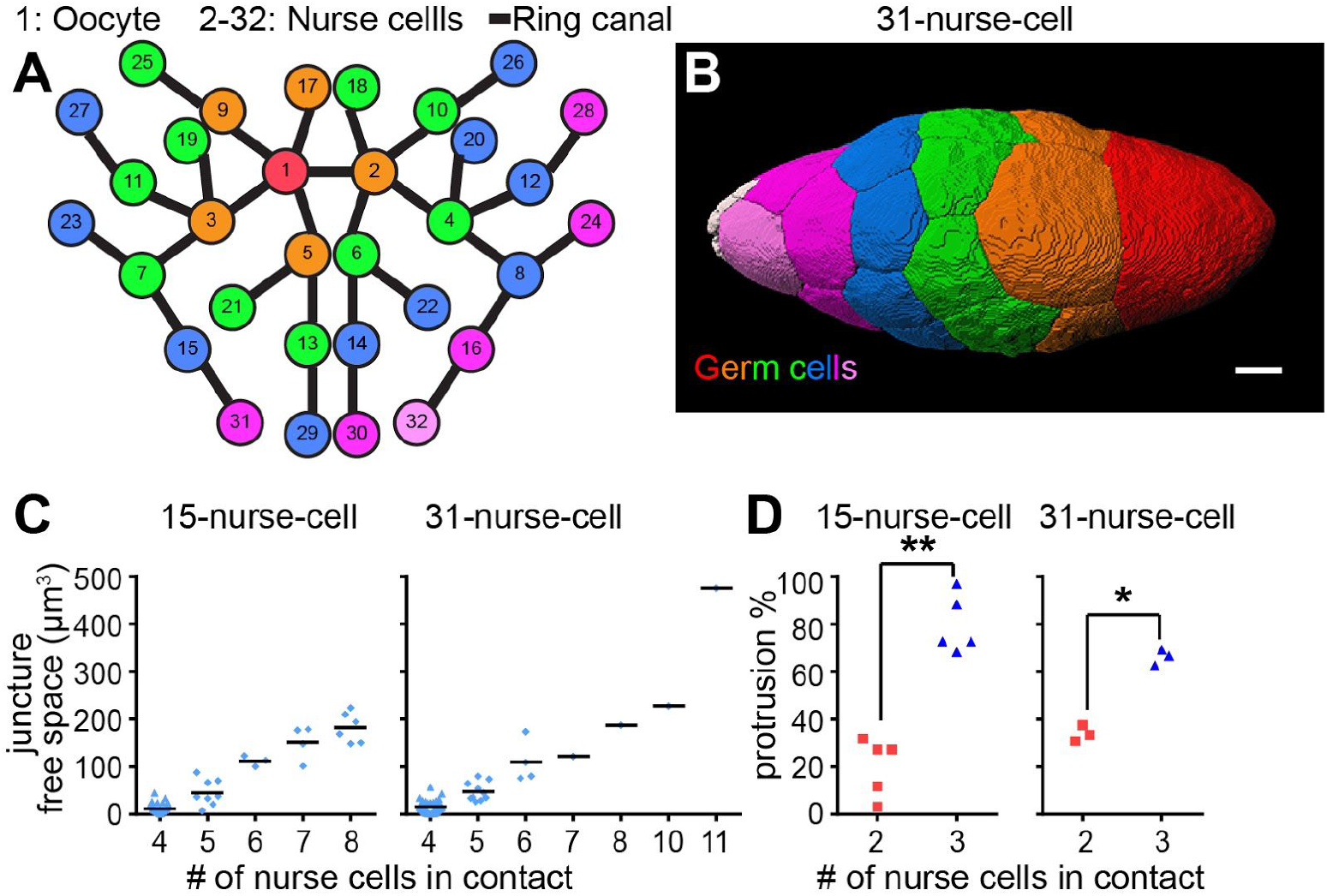
Nurse cell arrangements in mutants with increased number of nurse cells. (A) Schematic of the pattern of ring canal (lines) connections between individual germ cells (numbers in circles) in a 31-nurse-cell egg chamber. (B) 3D reconstruction of germ cell packing in a 31-nurse-cell stage 9 egg chamber. Germ cells are color coded according to ring canal number, which also correlates with proximity to the oocyte (red cell, #1). Border cells are at the anterior tip in B and are pseudocolored in white. Scale bar, 20 μm. (C) Quantification of extracellular spaces filled with fluorescent dextran in control and 31-nurse-cell egg chambers (n = 7, 3 egg chamber). (D) Quantification of side protrusion preferences as a fraction of total protrusions in control (n=5) and 31-nurse-cell (n=3) movies. **, *P* < 0.01, *, *P* < 0.05 (paired t test). Neither intercellular spaces nor protrusion preference for 3-nurse-cell junctures was altered in 31-nurse-cell egg chambers.

**Fig. S14.**
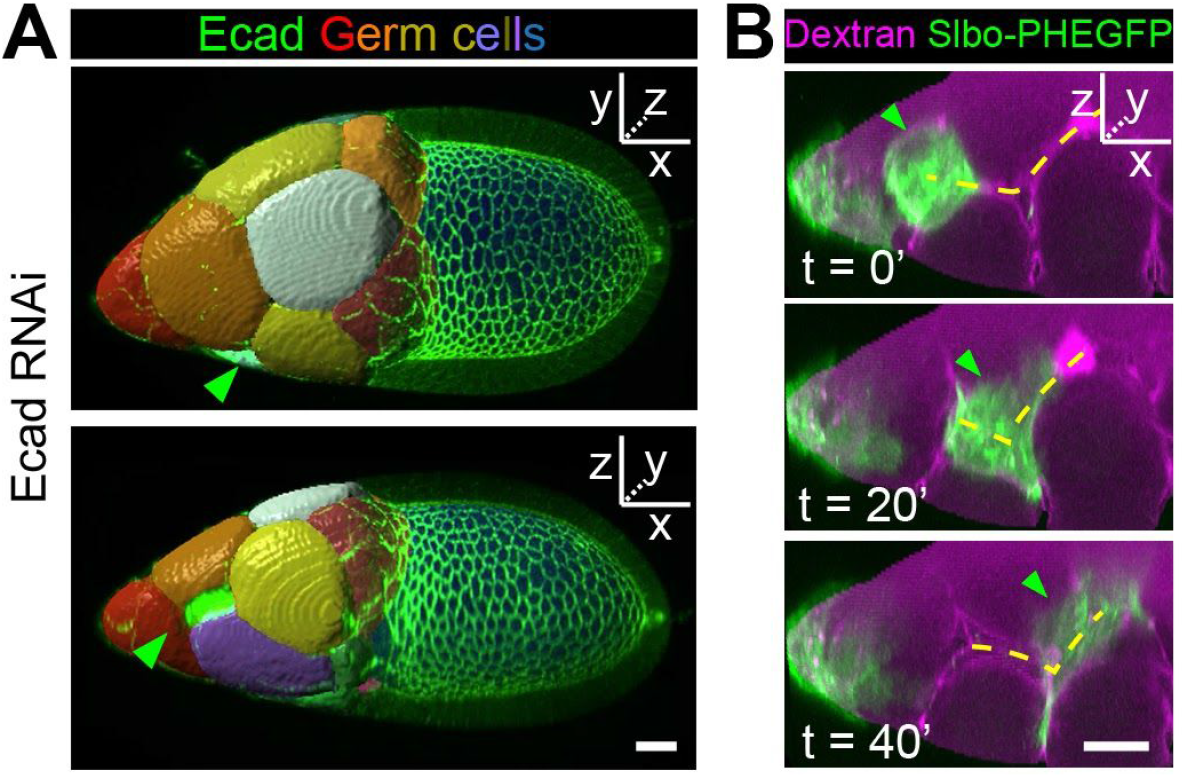
Border cells prefer multiple cell junctures in egg chambers with reduced germline Ecad. (A) Three dimensional reconstructions of nurse cells in a fixed egg chamber with germline Ecad RNAi showing border cells in nurse cell/nurse cell/follicle cell grooves. (B) Still images from a movie showing border cells (green arrowhead) zig zagging(dashed yellow lines) along grooves. Yellow dashed line indicates the nurse cell junctions that the border cells migrate along. Green arrowhead points to the border cells. Scale bars, 20 μm.

**Fig. S15.**
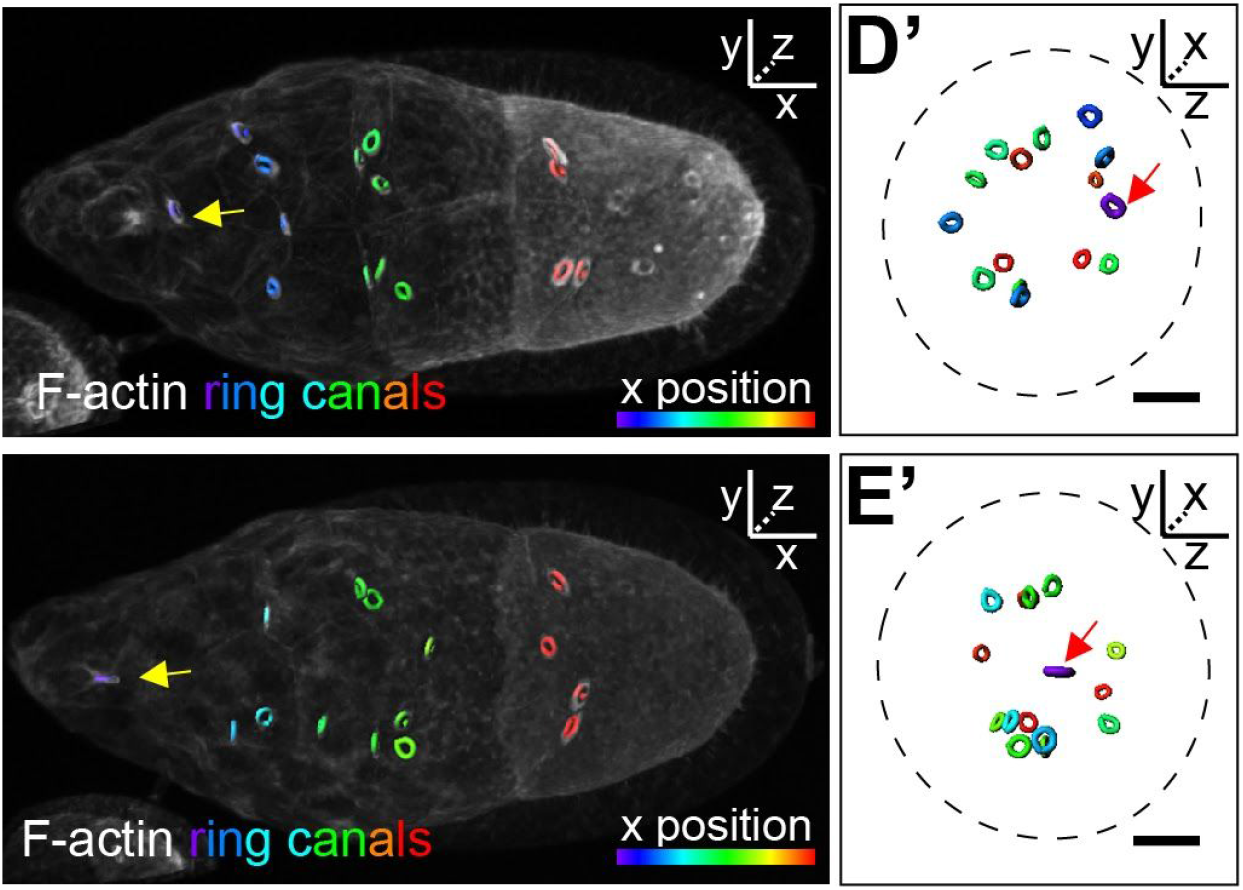
Ring canals are neither necessary nor sufficient to steer border cells. Lateral and anterior 3D projection views of reconstructed 3D ring canals (colored circles) in early stage 9 egg chambers in which ring canals are absent (upper right panel) or present (lower right panel) in the central path. Scale bars, 20 μm.

**Fig. S16.**
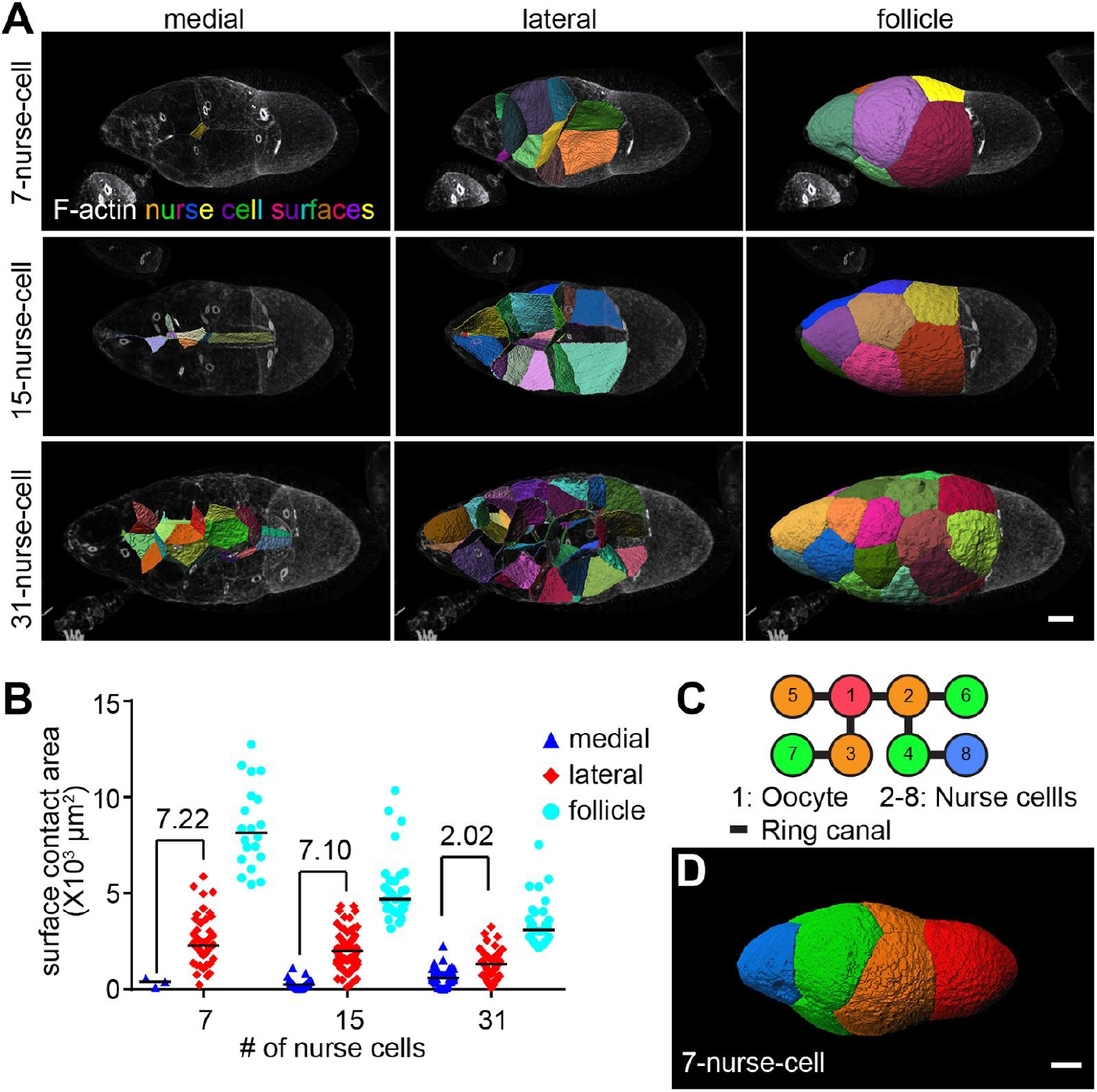
The ratio of medial to lateral surface areas is not critical for border cell migration. (A) Lateral view of segmented 3D nurse cell surface-surface contacts in early stage 9 egg chambers. (B) Quantification of the areas of different types of nurse cell surface contacts. Each dot represents the area of the contact between a single nurse cell and a neighboring nurse cell or follicle cells. Numbers indicate the mean value normalized to that of the medial path in each genotype to compare relative differences between medial and lateral surface areas. Data from *n* = 3, 2, 1 early stage 9 egg chambers. Absolute size of medial surfaces: 15NC (mean ± SD = 286 ± 256) and 31NC (mean ± SD = 658 ± 462) are significantly different (*P* < 0.0001, unpaired t test). 15NC and 7NC (mean ± SD = 347 ± 247) are not significantly different. Note there is only 1 medial surface in each 7NC egg chamber. (C) Schematic drawing of the pattern of ring canal (lines) connections between individual germ cells (numbers in circles) in a 7-nurse-cell egg chamber. (D) 3D reconstruction of germ cell packing in a 7-nurse-cell early stage 9 egg chambers. Germ cells are color coded according to ring canal number, which also correlates with proximity to the oocyte (red cell, #1). Scale bars, 20 μm.

### Supplementary Tables

**Table S1.**
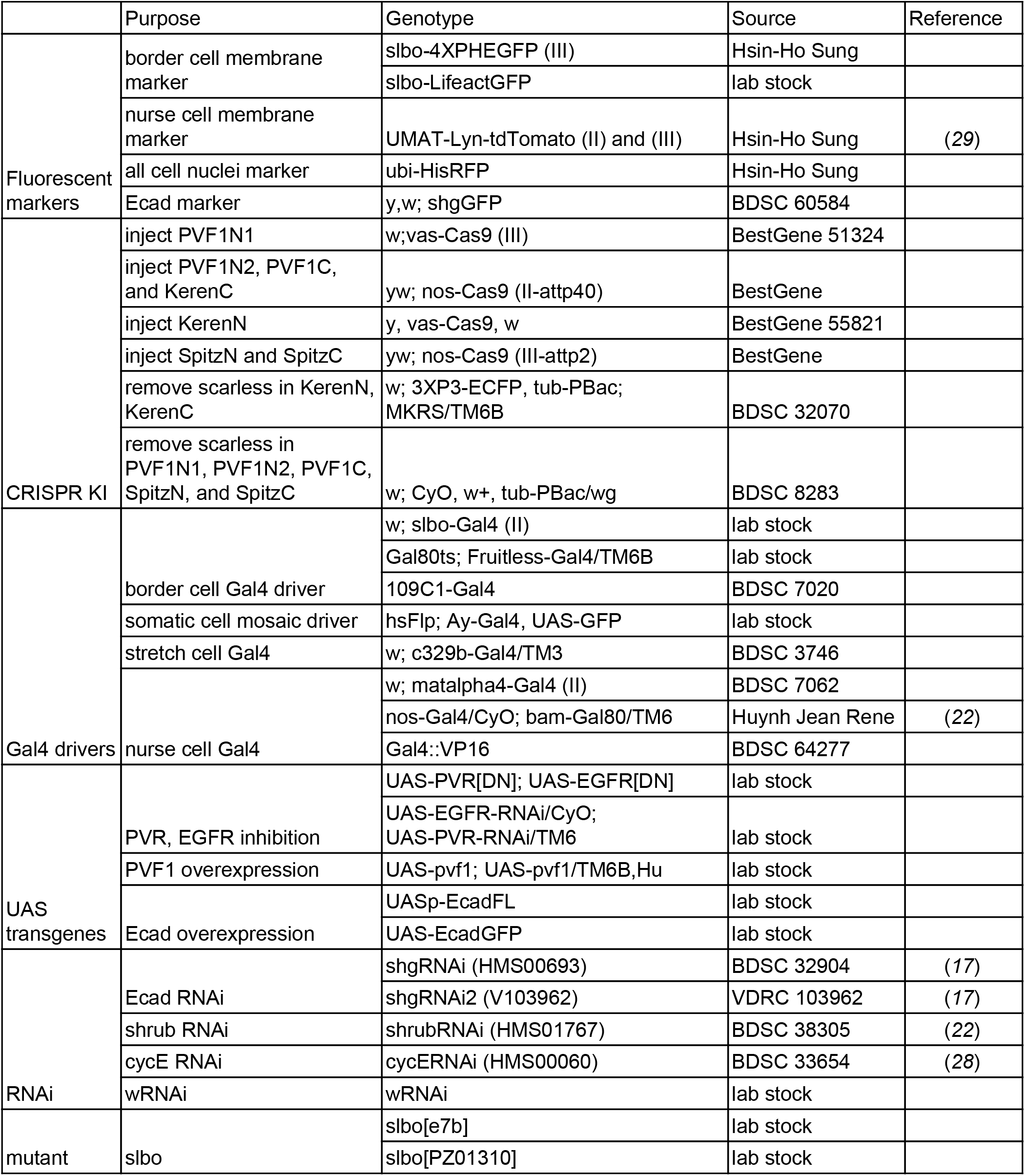
List of fly stains used in this study

**Table S2.**
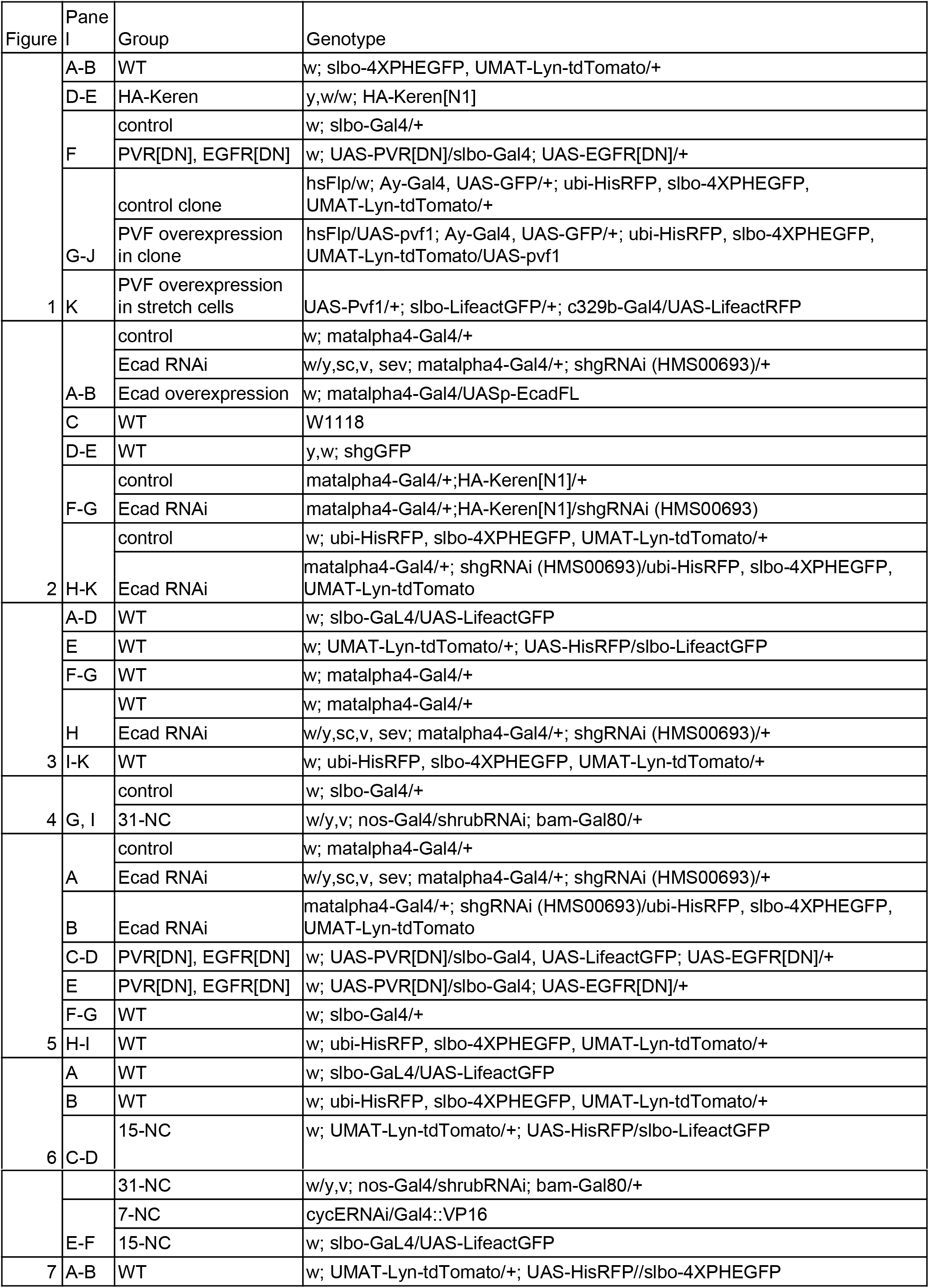
List of fly genotypes in each experiment

**Table S3.**
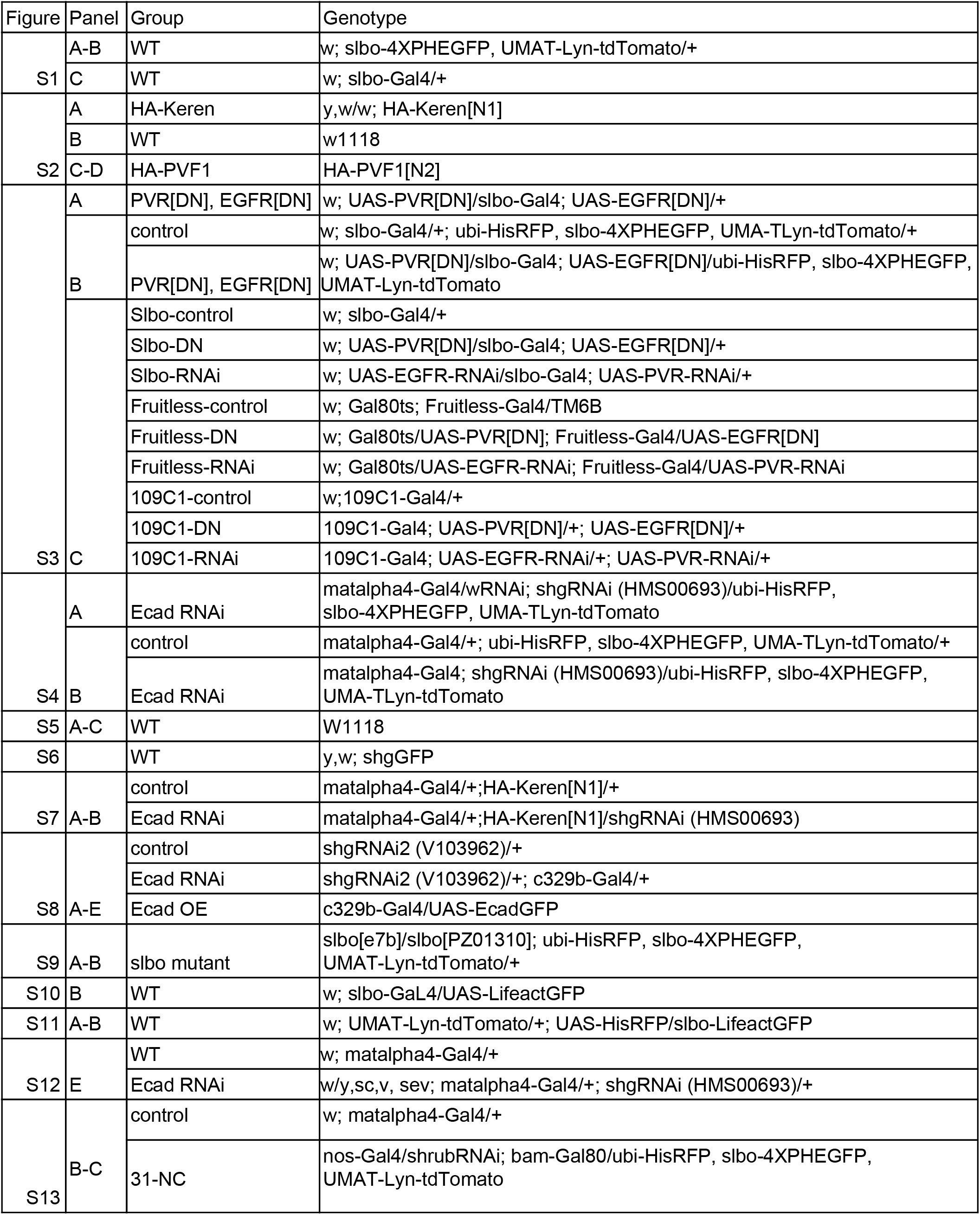
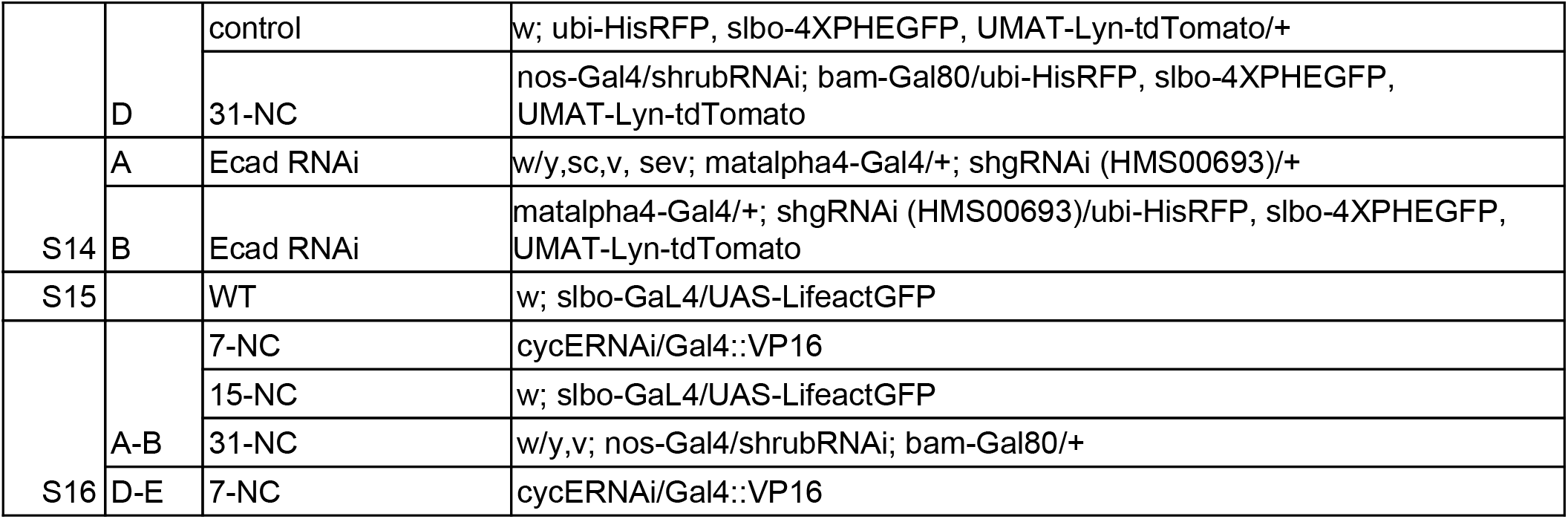
List of fly genotypes in each supplementary experiment

**Table S4.**
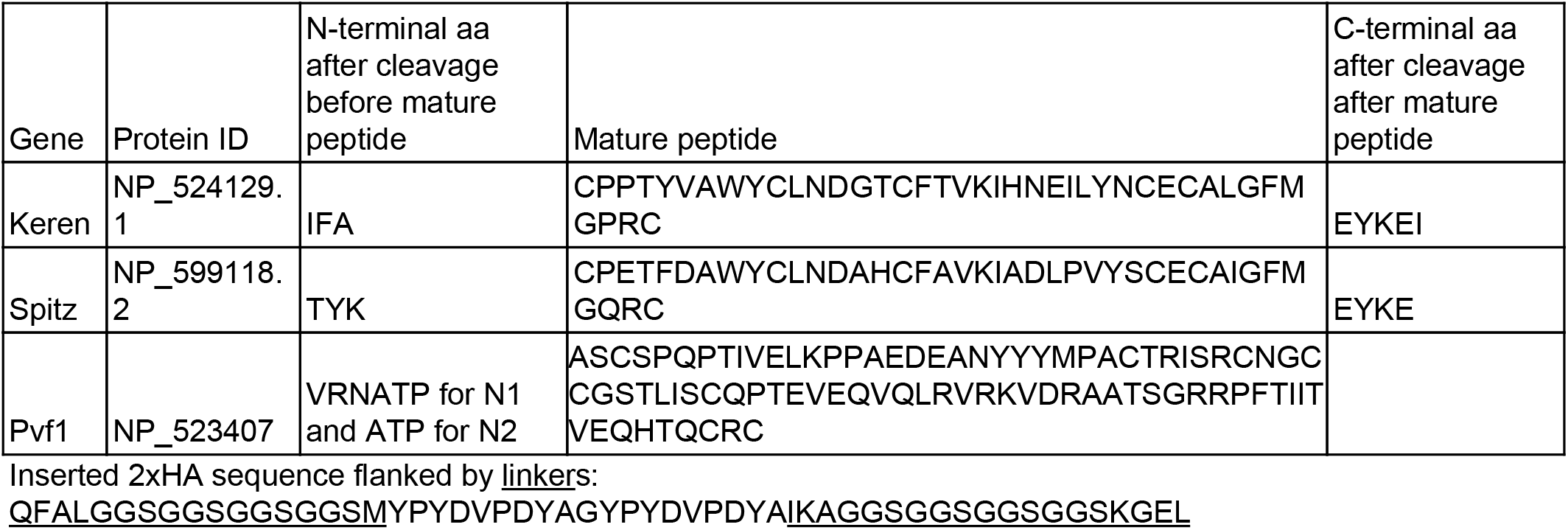
Predicted mature peptide and cleavage sites

**Table S5.**
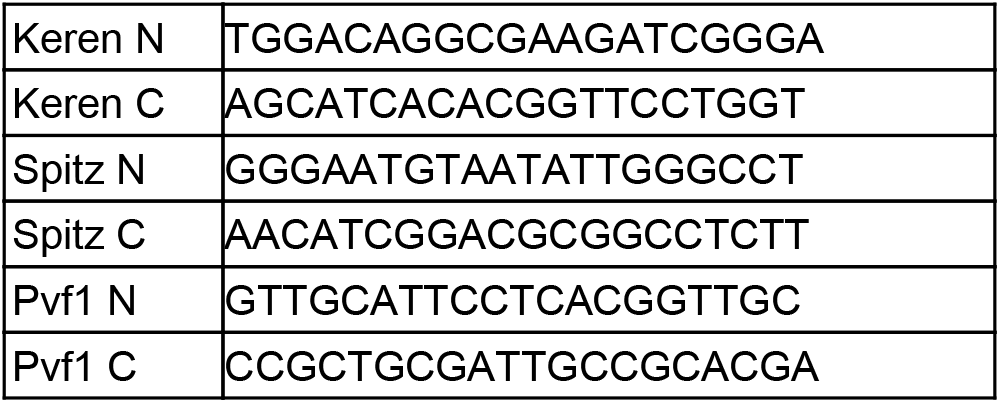
CRISPR target sites

**Table S6.**
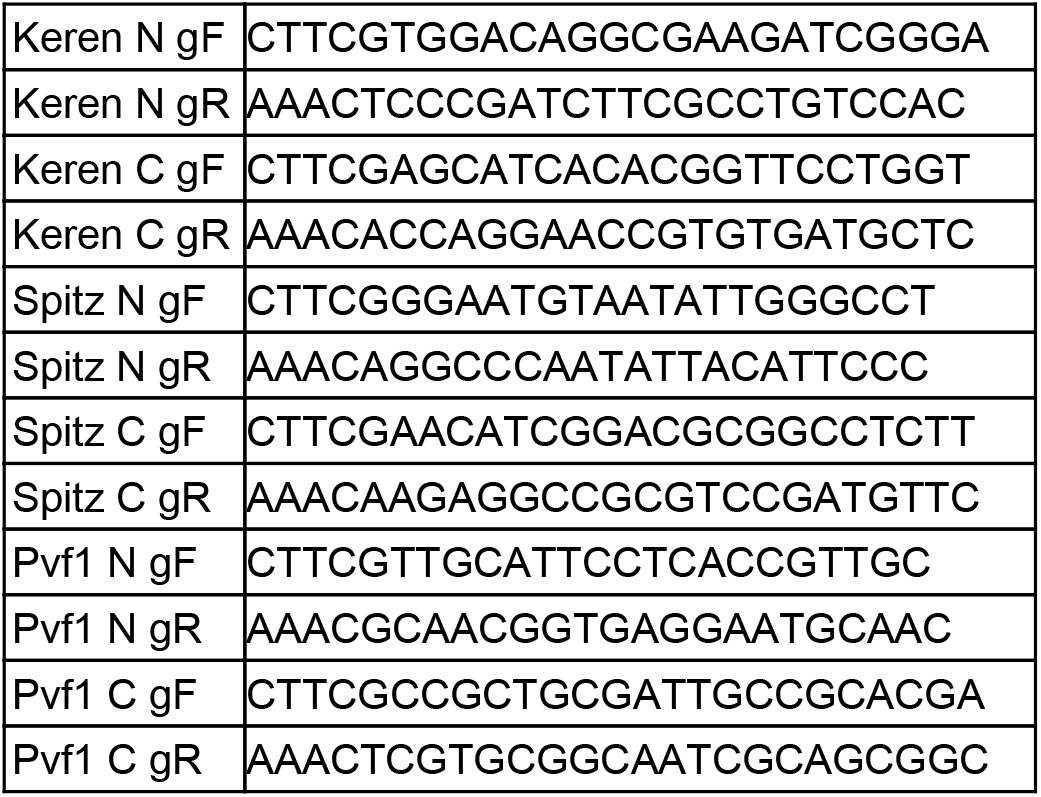
Primers for cloning CRISPR target into the pU6-BbsI-chiRNA vector

**Table S7.**
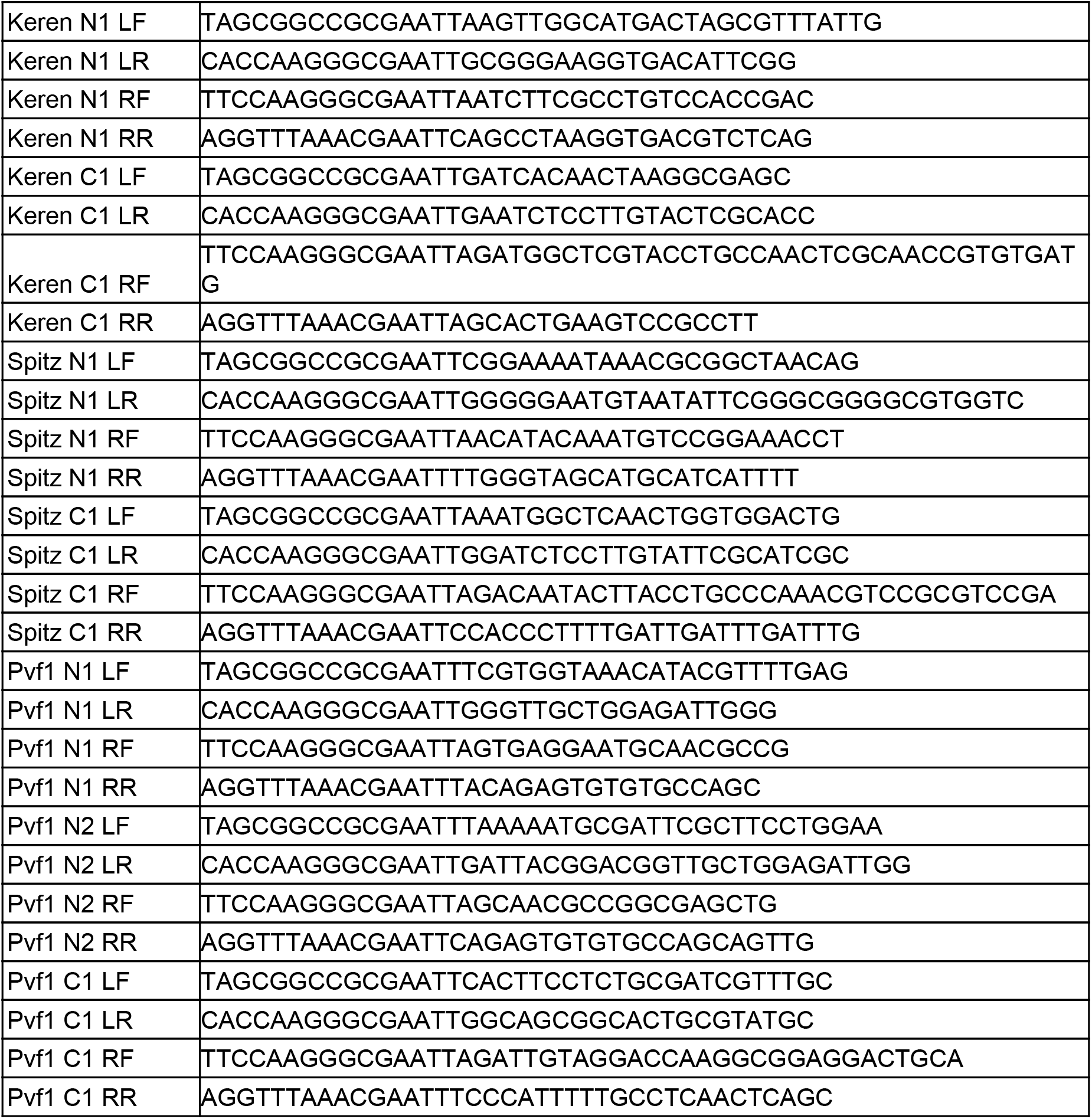
Primers for cloning left and right recombination arm into the pHD-2xHA-ScarlessDsRed vector.

**Table S8.** List of CRISPR knockin of HA tag in mature chemoattractant ligand peptide

|  | pU6-BbsI-chiRNA vector | pHD-2xHA-ScarlessD sRed vector | signal in live staining | signal in fixed staining | homozygous viable |
| --- | --- | --- | --- | --- | --- |
| Keren N1 | pU6 KerenN | pHD KerenN1 | yes | no | yes |
| Keren C1 | pU6 KerenC | pHD KerenC1 | no | no | no |
| Spitz N1 | pU6 SpitzN | pHD SpitzN1 | no | no | no |
| Spitz C1 | pU6 SpitzC | pHD SpitzC1 | no | no | no |
| PVF1 N1 | pU6 PVF1N | pHD PVF1N1 | no | N.A. | N.A. |
| PVF1 N2 | pU6 PVF1N | pHD PVF1N2 | no | yes | yes |
| PVF1 C1 | pU6 PVF1C | pHD PVF1C1 | no | no | no |

**Table S9.** Adhesion energy cost for N-cell junctures (in unit of the adhesion strength *A*)

| | $W_{adh,2}$ | $W_{adh,3}$ | $W_{adh,4}$ | $W_{adh,5}$ | $W_{adh,6}$ | $W_{adh,7}$ | $W_{adh,8}$ |
| --- | --- | --- | --- | --- | --- | --- | --- |
| value | $6.3 A$ | $4.9 A$ | $3.8 A$ | $2.8 A$ | $1.9 A$ | $1.1 A$ | $0.3 A$ |

**Table S10.** Model parameters.

| | $c_m$ | $L$ | $\xi$ | $r_0$ | $c_2$ | $\alpha$ | $\beta_1$ | $\beta_2$ |
| --- | --- | --- | --- | --- | --- | --- | --- | --- |
| value | 10 | 140 $\mu\text{m}$ | 42 $\mu\text{m}$ | 10 $\mu\text{m}$ | 1 | 80 | 0.4 | 0.2 |

**Table S11.** Parameter scanning of α and β_1_ Values in the table correspond to posterior and medial migration index and are color coded with red/green corresponding to low/high values. The parameter values used in wild-type simulations are underlined.

| % posterior | | $\beta_1$ | | | | | | | | | | |
| --- | --- | --- | --- | --- | --- | --- | --- | --- | --- | --- | --- | --- |
|  |  | 0 | 0.1 | 0.2 | 0.3 | 0.4 | 0.5 | 0.6 | 0.7 | 0.8 | 0.9 | 1 |
| $\alpha$ | 0 | 35 | 57 | 72 | 87 | 91 | 92 | 92 | 91 | 92 | 92 | 92 |
|  | 10 | 37 | 57 | 76 | 87 | 91 | 92 | 92 | 92 | 92 | 92 | 92 |
|  | 20 | 41 | 57 | 82 | 89 | 91 | 92 | 92 | 92 | 92 | 92 | 92 |
|  | 30 | 42 | 59 | 78 | 91 | 92 | 92 | 92 | 93 | 92 | 92 | 92 |
|  | 40 | 43 | 57 | 78 | 91 | 91 | 92 | 92 | 92 | 93 | 93 | 92 |
|  | 50 | 46 | 61 | 76 | 89 | 91 | 92 | 92 | 92 | 92 | 92 | 92 |
|  | 60 | 45 | 61 | 72 | 87 | 91 | 92 | 92 | 92 | 92 | 93 | 93 |
|  | 70 | 46 | 58 | 74 | 84 | 90 | 92 | 92 | 92 | 93 | 92 | 93 |
|  | 80 | 48 | 60 | 67 | 81 | <u>91</u> | 92 | 92 | 92 | 92 | 92 | 93 |
|  | 90 | 49 | 59 | 70 | 79 | 86 | 92 | 92 | 92 | 93 | 93 | 92 |
|  | 100 | 50 | 59 | 67 | 74 | 86 | 90 | 92 | 92 | 92 | 93 | 93 |
| % medial | | $\beta_1$ | | | | | | | | | | |
|  |  | 0 | 0.1 | 0.2 | 0.3 | 0.4 | 0.5 | 0.6 | 0.7 | 0.8 | 0.9 | 1 |
| $\alpha$ | 0 | 40 | 53 | 59 | 61 | 58 | 59 | 61 | 65 | 70 | 71 | 71 |
|  | 10 | 58 | 60 | 63 | 66 | 67 | 74 | 73 | 73 | 74 | 72 | 76 |
|  | 20 | 64 | 69 | 74 | 70 | 73 | 79 | 76 | 76 | 80 | 78 | 78 |
|  | 30 | 71 | 74 | 78 | 77 | 77 | 81 | 78 | 81 | 82 | 84 | 79 |
|  | 40 | 69 | 78 | 80 | 83 | 83 | 82 | 82 | 85 | 82 | 87 | 82 |
|  | 50 | 83 | 82 | 82 | 83 | 84 | 84 | 87 | 84 | 87 | 87 | 85 |
|  | 60 | 85 | 87 | 86 | 86 | 86 | 86 | 87 | 85 | 86 | 86 | 88 |
|  | 70 | 88 | 87 | 88 | 86 | 87 | 88 | 88 | 88 | 88 | 88 | 89 |
|  | 80 | 89 | 89 | 89 | 87 | <u>87</u> | 87 | 87 | 89 | 88 | 88 | 89 |
|  | 90 | 88 | 88 | 89 | 88 | 87 | 89 | 89 | 89 | 90 | 89 | 89 |
|  | 100 | 90 | 90 | 90 | 88 | 89 | 88 | 89 | 88 | 90 | 89 | 90 |

### Multimedia Files

**Movie 1.** Confocal time-lapse imaging of normal border cell migration. SlboGal4 drives UAS-mCD8-EGFP (green) and UAS-RFPnls (magenta).

**Movie 2.** “Fly-through” from the anterior tip of the egg chamber where the border cells are located to oocyte border, using near isotropic light sheet imaging showing the nurse cell membranes encountered by the border cells as they migrate. Multiple nurse cells meet in the center of the egg chamber where there is high membrane curvature. Border cells encounter ~40 side paths along the way. Green, E-cadherin. Magenta, F-actin. Scale bar, 20 μm.

**Movie 3.** Confocal time-lapse imaging of border cell migration with germline knockdown of E-cadherin. Matalpha4-Gal4;UAS-EcadRNAi. Green, Slbo-PHEGFP; magenta, UMAT-Lyn-tdTomato.

**Movie 4.** (Fig. 2H and I) (A) Confocal time-lapse imaging of a nurse cell juncture pulled by incoming border cell cluster in a control egg chamber. Green, Slbo-PHEGFP; magenta, UMAT-Lyn-tdTomato, Ubi-HisRFP. Note that the juncture shown by the yellow filament is just an example. Other junctures that the border cells contact are also deflected. In the first and last time points, tracking is removed to show the original membrane signal. (B) Confocal time-lapse imaging of an egg chamber with germline knockdown of E-cadherin (Matalpha4-Gal4;UAS-EcadRNAi) showing absence of nurse cell membrane deflection when border cell protrusions touch nurse cell membranes. Despite touching the nurse cell juncture multiple times, the border cell protrusion did not deflect it (one juncture is highlighted by the yellow filament). In the first and last time points, tracking is removed to show the original membrane signal.

**Movie 5.** (Fig. 3A and C) Animation of 3-dimensional reconstruction of different types of nurse cell contacts in a stage 9 egg chamber. Green, Slbo-LifeactGFP; white, F-actin or F-actin with outer follicle cell signal masked. Two-cell-contacts are shown as surfaces and color represents position along the z axis. Three-nurse cell junctures are represented as lines and color provides position in z. Magenta dots represent junctures of >3 nurse cells. First F-actin (white) and border cells (green) are shown. Then outer-follicle cell F-actin signals are masked to show nurse cell contacts. Then 2-cell surfaces are shown in 360 degree rotation, followed by >=3 cell juncture views in another 360 degree rotation.

**Movie 6.** (Fig. 4A) Simulation of border cell migration with nurse cell geometry and anterior-posterior chemoattractant gradient.

**Movie 7.** (Fig. 5B, fig. S14B). A confocal time-lapse movie of border cell migration in germline knockdown of E-cadherin. Border cells migrate along nurse-cell/nurse-cell/follicle-cell paths instead of nurse-cell/follicle-cell. Green, Slbo-PHEGFP; magenta, dextran.

**Movie 8.** (Fig. 6A, fig. S10B) Animation of 3-dimensional reconstruction of nurse cells and ring canals in a stage 9 egg chamber. White, F-actin; green, Slbo-LifeactGFP; blue, Hoechst. Nurse cell color indicates connection distance to the oocyte (red). White, border cells at the anterior tip. Yellow circles, ring canals. The egg chamber was stained to show F-actin (white), border cells (green), and nuclei (blue). First the reconstructed nurse cells rotate 360 degree to show that nurse cells that connect closer to the oocyte (red) are located more posteriorly. Then the ring canals (yellow) are shown to display their location in between connected nurse cells.

**Movie 9.** (Fig. 7A) A confocal time-lapse movie of wild-type border cell migration. Note the dorsal migration prior to reaching the oocyte boundary. Green, Slbo-PHEGFP; white, UMAT-Lyn-tdTomato.

**Movie 10.** (Fig. 7C) Simulation of border cell migration with nurse cell geometry and anterior-posterior chemoattractant gradient and dorsal Grk gradient.

## References

1. D. Cai, D. J. Montell, Diverse and dynamic sources and sinks in gradient formation and directed migration. Curr. Opin. Cell Biol. 30, 91–98 (2014).

2. Y. Artemenko, T. J. Lampert, P. N. Devreotes, Moving towards a paradigm: common mechanisms of chemotactic signaling in Dictyostelium and mammalian leukocytes. Cell. Mol. Life Sci. 71, 3711–3747 (2014).

3. K. F. Swaney, C.-H. Huang, P. N. Devreotes, Eukaryotic chemotaxis: a network of signaling pathways controls motility, directional sensing, and polarity. Annu. Rev. Biophys. 39, 265–289 (2010).

4. D. J. Montell, W. H. Yoon, M. Starz-Gaiano, Group choreography: mechanisms orchestrating the collective movement of border cells. Nat. Rev. Mol. Cell Biol. 13, 631–645 (2012).

5. E. Scarpa, R. Mayor, Collective cell migration in development. J. Cell Biol. 212, 143–155 (2016).

6. P. Friedl, D. Gilmour, Collective cell migration in morphogenesis, regeneration and cancer. Nat. Rev. Mol. Cell Biol. 10, 445–457 (2009).

7. P. Duchek, P. Rørth, Guidance of cell migration by EGF receptor signaling during Drosophila oogenesis. Science. 291, 131–133 (2001).

8. P. Duchek, K. Somogyi, G. Jékely, S. Beccari, P. Rørth, Guidance of cell migration by the Drosophila PDGF/VEGF receptor. Cell. 107, 17–26 (2001).

9. J. A. McDonald, E. M. Pinheiro, D. J. Montell, PVF1, a PDGF/VEGF homolog, is sufficient to guide border cells and interacts genetically with Taiman. Development. 130, 3469–3478 (2003).

10. J. A. McDonald, E. M. Pinheiro, L. Kadlec, T. Schupbach, D. J. Montell, Multiple EGFR ligands participate in guiding migrating border cells. Dev. Biol. 296, 94–103 (2006).

11. J. Bussmann, E. Raz, Chemokine-guided cell migration and motility in zebrafish development. EMBOJ. 34, 1309–1318 (2015).

12. R. Mayor, E. Theveneau, The neural crest. Development. 140, 2247–2251 (2013).

13. B. E. Richardson, R. Lehmann, Mechanisms guiding primordial germ cell migration: strategies from different organisms. Nat. Rev. Mol. Cell Biol. 11, 37–49 (2010).

14. M. R. Ng, A. Besser, G. Danuser, J. S. Brugge, Substrate stiffness regulates cadherin-dependent collective migration through myosin-II contractility. J. Cell Biol. 199, 545–563 (2012).

15. G. Aranjuez, A. Burtscher, K. Sawant, P. Majumder, J. A. McDonald, Dynamic myosin activation promotes collective morphology and migration by locally balancing oppositional forces from surrounding tissue. Mol. Biol. Cell. 27, 1898–1910 (2016).

16. E. H. Barriga, K. Franze, G. Charras, R. Mayor, Tissue stiffening coordinates morphogenesis by triggering collective cell migration *in vivo*. Nature. 554, 523–527 (2018).

17. D. Cai et al., Mechanical feedback through E-cadherin promotes direction sensing during collective cell migration. Cell. 157, 11461159 (2014).

18. J. I. Alsous, P. Villoutreix, N. Stoop, S. Y. Shvartsman, J. Dunkel, Entropic effects in cell lineage tree packings. Nat. Phys. 14, 10161021 (2018).

19. M. R. Zanotelli et al., Energetic costs regulated by cell mechanics and confinement are predictive of migration path during decisionmaking. Nat. Commun. 10, 4185 (2019).

20. J. Renkawitz et al., Nuclear positioning facilitates amoeboid migration along the path of least resistance. Nature. 568, 546–550 (2019).

21. N. R. Matias, J. Mathieu, J.-R. Huynh, Abscission is regulated by the ESCRT-III protein shrub in Drosophila germline stem cells. PLoS Genet. 11, e1004653 (2015).

22. M. A. Lilly, A. C. Spradling, The Drosophila endocycle is controlled by Cyclin E and lacks a checkpoint ensuring S-phase completion. Genes Dev. 10, 2514–2526 (1996).

23. X. Wang, L. He, Y. I. Wu, K. M. Hahn, D. J. Montell, Light-mediated activation reveals a key role for Rac in collective guidance of cell movement *in vivo*. Nat. Cell Biol. 12, 591–597 (2010).

24. W. Dai, D. J. Montell, Live imaging of border cell migration in drosophila. Methods Mol. Biol. 1407, 153–168 (2016).

25. S. Preibisch et al., Efficient Bayesian-based multiview deconvolution. Nat. Methods. 11, 645–648 (2014).

26. I. Heemskerk, S. J. Streichan, Tissue cartography: compressing bio-image data by dimensional reduction. Nat. Methods. 12, 11391142 (2015).

27. D. Cai et al., Modeling and analysis of collective cell migration in an *in vivo* three-dimensional environment. Proc Natl Acad Sci USA. 113, E2134–41 (2016).

28. D. J. Montell, P. Rorth, A. C. Spradling, slow border cells, a locus required for a developmentally regulated cell migration during oogenesis, encodes Drosophila C/EBP. Cell. 71, 51–62 (1992).

29. A. Cliffe et al., Quantitative 3D analysis of complex single border cell behaviors in coordinated collective cell migration. Nat. Commun. 8, 14905 (2017).

